# Decoding the *MYC* locus reveals a druggable ultraconserved RNA element

**DOI:** 10.64898/2026.01.29.702547

**Authors:** Peiguo Shi, Feiyue Yang, FNU Tala, Wesley Huang, Alexis O. Aparicio, Colin H. Kalicki, Aditi Trehan, Michael R. Murphy, Esther R. Rotlevi, Linqing Xing, Muredach P. Reilly, Jianwen Que, Xuebing Wu

**Affiliations:** Division of Cardiology, Department of Medicine, Vagelos College of Physicians and Surgeons, Columbia University Irving Medical Center, New York, NY 10032, USA; Department of Systems Biology, Vagelos College of Physicians and Surgeons, Columbia University Irving Medical Center, New York, NY 10032, USA; Herbert Irving Comprehensive Cancer Center, Vagelos College of Physicians and Surgeons, Columbia University Irving Medical Center, New York, NY 10032, USA; Columbia Center for Human Development, Department of Medicine, Vagelos College of Physicians and Surgeons, Columbia University Irving Medical Center, New York, NY 10032, USA; Current address: Renaissance School of Medicine at Stony Brook University Stony Brook, Stony Brook, NY 11794, USA; Division of Digestive and Liver Disease, Department of Medicine, Vagelos College of Physicians and Surgeons, Columbia University Irving Medical Center, New York, NY 10032, USA

**Author notes:** Correspondence to (P.S.); (X.W.). Equal contribution (Peiguo Shi, Feiyue Yang).

## Abstract

The human genome is dominated by noncoding sequences, most of which are poorly conserved across species. How genetic information is distributed between coding and noncoding regions remains a fundamental unresolved question. Using CRISPR saturation mutagenesis at base-pair resolution, we mapped the functional fitness landscape of the 10-kb human *MYC* locus with a near-PAMless, high-fidelity SpRY-Cas9. This unbiased interrogation revealed that the majority (67%) of functionally essential base-pairs in this locus are noncoding. Paradoxically, the phenotypic impact of noncoding sequences correlates inversely with evolutionary conservation, driven in part by rapidly diverging cis-regulatory DNA elements that remain functionally constrained in humans. Within this landscape, we identified an ultraconserved RNA element in the 3’ untranslated region (UTR) that is indispensable for MYC-dependent cancer cells. Remarkably, steric-blocking antisense oligos targeting this RNA element selectively eliminate MYC-addicted cancer cells by suppressing MYC function without reducing MYC abundance. Mechanistically, this 3′ UTR element promotes perinuclear localization of *MYC* mRNA and efficient nuclear import of the short-lived MYC protein, enabling its function as a nuclear transcription factor. Together, these findings highlight noncoding sequences as major carriers of functional genetic information, provide a comprehensive fitness map of the *MYC* locus, and uncover a therapeutically actionable RNA element that disables MYC-driven cancer.

## INTRODUCTION

The vast noncoding expanse of the human genome represents one of the greatest remaining frontiers in understanding genome function^1^. Once dismissed as inert, noncoding sequences are now known to orchestrate gene expression at multiple layers – serving as promoters, enhancers and silencers at the DNA level^2^, and governing splicing, polyadenylation, subcellular localization, translation, and decay at the RNA level^3^. Yet despite these advances, the functional landscape of the noncoding genome remains poorly understood. Strikingly, although individual noncoding elements often show limited conservation, more than 70% of the base-pairs conserved between humans and mice are noncoding^4^. This observation implies that the majority of the genome’s genetic information may be embedded outside of coding regions. However, systematic and unbiased experimental evidence supporting functional contributions on this scale is still lacking, and the biological roles of most conserved noncoding elements remain unknown.

In contrast to coding sequences – whose functions and mutational impacts are relatively straightforward to predict and interrogate – the functions of noncoding regions are far more challenging to dissect, owing to the diverse mechanisms through which they act. Massively parallel reporter assays (MPRAs)^5^ have been widely used to test the activity of individual elements – such as putative enhancers^6^ and RNA elements regulating splicing^7^, polyadenylation^8^, stability^9^ or translation^10^ – in artificial reporter constructs. Although MPRAs enable the simultaneous testing of thousands of putative regulatory sequences and their variants, they lack the native genomic and transcript context and are often confounded by technical artifacts^11,12^. Alternatively, CRISPR-mediated saturating mutagenesis^13^ and CRISPRi-mediated tiling scan^14^ have been applied to identify functional elements within their endogenous genomic context^15^. However, the comprehensiveness and resolution of these studies have been limited, in part due to the dependence of SpCas9 on the canonical Protospacer Adjacent Motif (PAM) NGG, which restricts the density of targetable sites to roughly every 8-12 bps.

MYC is a master regulator of cell proliferation and one of the most frequently dysregulated oncogenes, with aberrant expression observed in approximately 70% of human cancers^16,17^. Many tumors exhibit “MYC addiction,” relying on sustained MYC activity for proliferation and survival^18,19^. Despite decades of research, no approved therapy directly targeting the MYC protein exists^20^, due in part to its intrinsically disordered structure^20^, predominant nuclear localization^21^, and an incomplete understanding of its biology. A deeper characterization of MYC’s regulation may therefore open new therapeutic avenues.

Using the *MYC* locus as a proof of principle, we conducted, to our knowledge, the first base-pair-resolution saturated CRISPR mutagenesis screen of an entire protein-coding gene, enabled by a near-PAMless high-fidelity Cas9 variant (HF-SpRY-Cas9)^22^. Our unbiased screen revealed the fitness landscape of the *MYC* locus and uncovered a previously hidden 3’ UTR regulatory element essential for MYC’s oncogenic activity. We further developed antisense oligonucleotides (ASOs) targeting this ultraconserved element, which potently suppress MYC-dependent cancer cell growth. Our study establishes a generalizable platform for base-pair-resolution functional dissection of endogenous genes and provides a powerful strategy for therapeutic discovery.

## RESULTS

### CRISPR saturation mutagenesis screening of the *MYC* locus

To comprehensively assess the phenotypic impact of each base-pair (bp) within the 10,055-bp human *MYC* locus (Fig. 1A), we performed a cell growth-based saturation mutagenesis screen using HF-SpRY-Cas9 and a library of sgRNAs tiling both strands of the locus. Taking advantage of the near-PAMless activity of SpRY-Cas9, we designed sgRNAs for all possible 20-nt sequences, enabling each base-pair to be interrogated by four sgRNAs that are expected to induce double-strand breaks immediately flanking it (red sgRNAs targeting the blue base-pair in Fig. 1B). In total, the library comprised 22,623 sgRNAs, including 565 negative controls.

**Figure 1:**
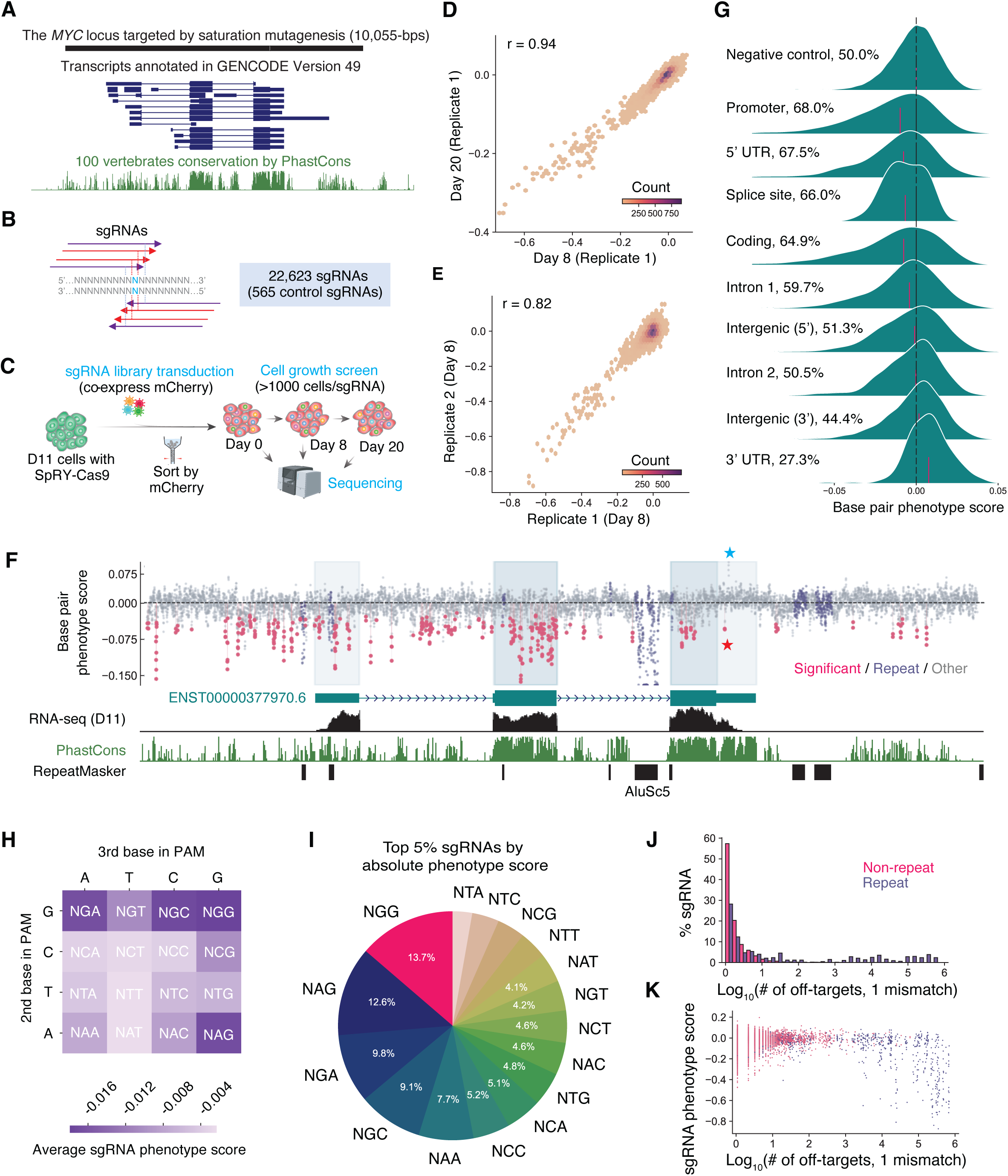
Mapping the fitness landscape of the *MYC* locus with CRISPR saturation mutagenesis. **A**. The *MYC* locus targeted by CRISPR mutagenesis, together with annotated *MYC* transcripts and phastCons conservation score across 100 vertebrate genomes. **B**. Each base-pair (light blue) is targeted by four sgRNAs (red arrows), two on each strand, that induce Cas9 cleavage (dotted vertical lines) on either side. Four additional sgRNAs (purple arrows) induce cleavage within ±1 bp. **C**. Schematic of the CRISPR saturation mutagenesis screen for cell proliferation. **D**. Scatter plot of base-pair level phenotype scores between day 8 and day 20 samples (replicate 1). **E**. Scatter plot of base-pair level phenotype scores between replicate 1 and replicate 2 (day 8 samples). **F**. Visualization of the base-pair level phenotype scores averaged across four samples. Significant base-pairs (methods specified in Methods: base-pair phenotype score calculation) are highlighted in red. Repeats are highlighted in purple. Scores are truncated at the AluSc5 repeat (see Fig. S2 for the full range). Blue and red stars highlight 3’ UTR base pairs with the strongest positive and negative phenotype scores, respectively. **G**. Density plots of base-pair phenotype scores in various regions. The vertical red bar indicates median values. Also shown are the percentage of base-pairs with a negative phenotype score. Promoter is defined as the 1-kb region upstream of the transcription start site of transcript ENST00000377970.6. **H**. Average phenotype scores across four samples of all sgRNAs with the same PAM. A dark color indicates a strong negative phenotype (deleterious). **I**. Frequency of PAMs associated with top 5% sgRNAs by absolute sgRNA-level phenotype scores average across four samples. **J**. Distribution of sgRNAs binned by the number of off-targets with one mismatch. **K**. Scatter plot of sgRNA phenotype scores across four samples and off-target counts (allowing one mismatch).

The screen was conducted in the MYC-dependent human D11 cell line, a derivative of the multiple myeloma cell line JJN-3 that lacks *MYC* gene amplification^23,24^ and requires MYC for proliferation^25^. In D11 cells, one *MYC* allele carries a C-terminal 2A-d2GFP knockin, enabling fluorescence-based monitoring of *MYC* expression using the short-lived d2GFP^24^. We first generated a D11 cell line stably expressing a high-fidelity SpRY-Cas9 variant through lentiviral transduction followed by puromycin selection (Fig. 1C). Cells were subsequently transduced with the sgRNA library at a low multiplicity of infection (MOI<0.3), selected by mCherry sorting (co-express with sgRNAs), and cultured while maintaining >1000 cells per sgRNA. Cells were harvested on day 0, 8, and 20 for sgRNA sequencing. An independent biological replicate was performed starting from SpRY-Cas9 stable cell line generation.

Using MAGeCK^26^, we calculated a phenotype score for each sgRNA as the log_2_ fold-change in abundance at day 8 or day 20 relative to day 0, normalized by negative control sgRNAs (Supplementary Table 1). To experimentally validate the screen, we generated cell lines each stably expressing one of 12 strongly depleted sgRNAs and validated the presence of substantial indels at all target sites using amplicon sequencing (Fig. S1A-B, related oligos in Supplementary Table 2). Six of these cell lines were further evaluated in competitive growth assay, and all consistently reduced cell fitness compared with negative control (Fig. S1C), confirming the robustness of this screening system.

### Reproducible phenotypes of every base-pair

To quantify phenotypic effect at base-pair resolution, we applied MAGeCK’s Robust Rank Aggregation (RPA) algorithm^26^ to integrate data from the eight sgRNAs predicted to cut within ±1 bp of each base-pair (Fig. 1B), as these guides are most likely to induce indels at that position. Aggregating eight sgRNAs per base-pair reduces function-independent variability arising from differences in sgRNA efficiency, PAM compatibility, and off-target effects. Base-pair-level phenotype scores (Supplementary Table 3) were highly correlated between day 8 and day 20 samples within each replicate (Pearson correlation coefficient *r* = 0.94 for replicate 1, Fig. 1D, *r* = 0.86 for replicate 2, Fig. S1D). Across replicates, correlations remained strong, with *r* = 0.82 for day 8 samples (Fig. 1E) and *r* = 0.74 for day 20 samples (Fig. S1E). These results demonstrate the high reproducibility of our screening approach. Given the strong agreement between replicates and time points, we combined all four samples (two replicates each with two time points) and calculated a composite phenotype score for each base-pair by averaging across samples. This analysis delineates the fitness landscape of the *MYC* locus at base-pair resolution (Fig. 1F). A base-pair position was considered to have a significant phenotype if it showed an FDR < 0.05 in at least two samples and exhibited a consistent directional trend in the remaining samples (N=343, highlighted in red in Fig. 1F).

Multiple lines of evidence indicate that our screen performed as expected. First, the strongest phenotype is mapped to an intronic *Alu* repeat element (Fig. S2A), which is found with over one million copies in the genome and CRISPR targeting of such high-copy sites is known to cause extensive DNA double-strand breaks and severely impair cell viability non-specifically^27^. Deletion of the Alu element from an 8.6-kb fragment encoding the full *MYC* gene on a plasmid had no detectable effect on *MYC* mRNA splicing or abundance, as assessed by RNA-seq (Fig. S2C). All repeat regions (annotated in *RepeatMasker*) were excluded from subsequent analysis unless otherwise noted (colored purple in Fig. 1F). Second, among non-repeat regions, we observed strong growth defects (negative phenotype scores) associated with the promoter, 5’ UTR, splice sites, and coding region (Fig. 1G), consistent with their essential role in MYC function. In contrast, targeting the 3’ UTR tends to promote cell growth (positive phenotype scores, Fig. 1G), consistent with *MYC* 3’ UTR’s role in promoting rapid RNA degradation^28^. Third, we observed a PAM preference similar to that previously described for SpRY-Cas9^10^, which displays the highest activity at NGN PAMs, with close to background activity for NTT and NAT PAMs (Fig. 1H). Notably, among the top 5% non-repeat sgRNAs with the strongest phenotype, 86.3% do not use the canonical NGG PAM, and thus would have been missed by a screen using the canonical SpCas9 (Fig. 1I). Lastly, we evaluated sgRNA activity using three widely used on-target efficiency prediction algorithms^29–31^ and observed significant correlation between predicted on-target efficiency and observed phenotype scores (Fig. S3).

### Negligible false positives linked to off-target effects

The HF-SpRY-Cas9 system used in this study is a high-fidelity Cas9 variant previously shown to have negligible off-target activity^22^. The majority (57%) of non-repeat-targeting sgRNAs in our library have no predicted off-target sites in the genome when allowing one mismatch (Fig. 1J). In contrast to repeat-targeting sgRNAs, which tend to harbor extremely large numbers of predicted off-target sites (∼10^3^ to ∼10^6^) and whose deleterious effects scale with off-target counts (Fig. 1K, purple), non-repeat-targeting sgRNAs generally have few predicted off-target sites and the phenotypic impact diminishes (scores approaching zero) as off-target counts increase (red, Fig. 1K). This trend is likely explained by dilution and sequestration of Cas9/sgRNA complexes at predominantly nonessential off-target loci, together with reduced cleavage efficiency at mismatched sites. These analyses indicate that off-target activity, if any, is more likely to dampen rather than enhance phenotypes, thereby contributing minimally to false positives for non-repeat-targeting sgRNAs.

Moreover, since the phenotypic score for each base-pair is aggregated across eight distinct sgRNAs (Fig. 1B), the influence of any off-target event associated with a single sgRNA is further minimized, including potential off-target effects caused by mutated sgRNA as a result of self-targeting^32^. Therefore, in subsequent analyses restricted to non-repeat sgRNAs, strong phenotypes are unlikely to arise from off-target effects. Nevertheless, in the mechanistic studies that follow, we systematically ruled out potential off-target contributions from individual sgRNAs using targeted amplicon sequencing. Lastly, the observed cell growth phenotype is unlikely to result from disruption of a neighboring gene, as the nearest protein-coding gene lies more than 300-kb away.

### The majority of functional base-pairs are noncoding

As expected, mutations in coding regions were more likely to produce significant phenotypic effects: although coding sequences comprise only 14% of the *MYC* locus interrogated (excluding repeats), they accounted for 33% of the base-pairs with significant phenotypic impact (Fig. 2A), representing a ∼3-fold enrichment relative to noncoding sequences. While this enrichment underscores the importance of individual coding base pairs, it paradoxically highlights that noncoding regions collectively harbor the majority (67%) of functional sites (Fig. 2A). Similar trends were observed when considering the top 5% base-pairs ranked by absolute phenotypic score (Fig. 2A).

**Figure 2.**
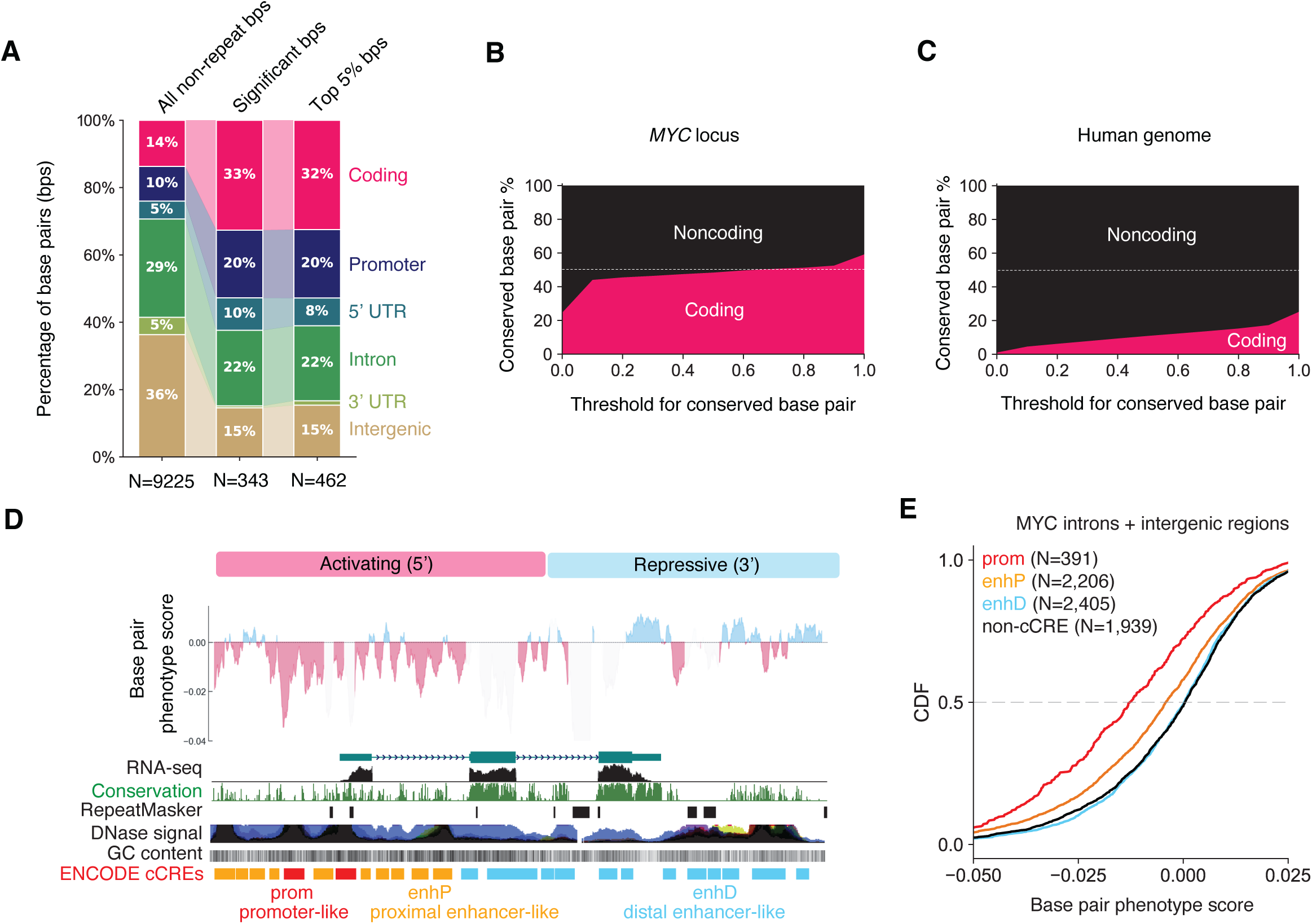
Noncoding region accounts for the majority of functional base-pairs in the *MYC* locus. **A**. Distribution of base-pairs (bps) in *MYC*, comparing those with significant phenotypes or within the top 5% by absolute phenotype scores to all non-repeat bps in the *MYC* locus. **B**. Relative percentage of coding and noncoding base-pairs in *MYC* gene (ENST00000377970.6) with conservation scores higher than a given phastCons score threshold. **C**. Same as B for the entire human genome. **D**. Phenotype scores of noncoding base-pairs smoothed using a sliding window of 100-bps. Signals in coding regions (except the end of the CDS containing an RNA instability element) and repeats are masked to highlight noncoding regions. ENCODE DNase I signal range: 0-100. **E**. Cumulative distribution function (CDF) plot comparing phenotype scores of base-pairs within ENCODE3 annotated cCREs (prom vs non-cCRE: P = 1.2x10^-19^, enhP vs non-cCRE: P = 2.6x10^-10^). Kolmogorov–Smirnov test was used to compute the P values.

These results were consistent with patterns of evolutionary conservation across 100 vertebrate species: across a wide range of conservation score thresholds, approximately 50% of the conserved base-pairs within the *MYC* locus are noncoding (Fig. 2B). Genome-wide, more than 70% of the most highly conserved base-pairs are noncoding (Fig. 2C), consistent with a previous estimate based on human-mouse sequence conservation^4^. These findings emphasize that although individual noncoding base-pairs are, on average, less likely to be functional, the majority of the genome’s functional genetic information may be collectively encoded within noncoding sequences.

Intriguingly, noncoding elements at the 5’ and 3’ halves of the locus exhibit opposing phenotypic impact, as revealed by phenotype scores smoothed using a sliding window of 100-bp, excluding coding regions (Fig. 2D). Splitting in intron 2, the 5’ half of the locus contains base-pairs that confer slower cell growth when mutated, including the majority (84%) of noncoding base-pairs with significant phenotypic effects (Fig. 1F), indicating a key role of the 5’ half in promoting MYC expression and function. In contrast, noncoding base-pairs in the 3’ half are mostly repressive, whose mutation mostly resulted in faster cell growth (Fig. 2D, and Fig. 1G), although none are significant (Fig. 1F).

The majority (79%) of significant 5’ noncoding base-pairs overlap with ENCODE-annotated candidate *cis*-regulatory elements (cCREs)^33^ bearing promoter-like (prom, 24%) or proximal enhancer-like (enhP, 54%) chromatin signatures (Fig. 2D), suggesting that these sequences regulate *MYC* transcription. Consistent with this interpretation, base-pairs within prom and enhP regions were significantly more likely to reduce cell viability in our CRISPR screen (Fig. 2E). In contrast, distal enhancer like signatures, mostly located in the 3’ half of the *MYC* locus (Fig. 2D), do not appear to be critical for the cell line we investigated (Fig. 2E). Together, these findings indicate that the 5’ half of the *MYC* locus functions as a transcriptional activation hub that drives *MYC* expression.

The majority of growth-suppressing sequences (positive phenotype scores) cluster in the 3’ half of the locus, particularly within the highly conserved, AU-rich 3’ UTR that mediates rapid *MYC* mRNA degradation^28^ (Fig. 2D), as well as the C-terminal coding region that induces ribosome stalling and mRNA decay^34^. However, no individual positions reach statistical significance (Fig. 1F). This includes the site with the strongest positive phenotype score across the entire locus (blue star, Fig. 1F, lowest FDR = 0.07), which notably resides within the most conserved region of the 3’ UTR – the only 9-nt or longer sequence (AACTGCCTC) perfectly conserved across at least 82 vertebrate genomes (ultraconserved, Fig. S4A). Intriguingly, this 9-nt element contains two overlapping seed-match sites, offset by a single nucleotide, for the highly conserved tumor suppressor microRNAs let-7b and miR-34b (Fig. S4B), both promoting *MYC* mRNA degradation^35–37^. Nonetheless, transducing individual sgRNAs targeting this miRNA site into SpRY-Cas9 D11 cells failed to generate significant indels, consistent with the lack of statistical significance in the screen. The generally weak growth-suppressive phenotypes observed across the 3’ UTR may reflect technical constraints of CRISPR targeting in this region, including markedly reduced chromatin accessibility (as indicated by low DNase signal in Fig. 2D) and a high AT content enriching for less efficient PAMs (Fig. 1H).

### A paradox between conservation and function

The *MYC* locus contains numerous evolutionarily conserved noncoding sequences (Fig. 1F). According to the prevailing view that sequence conservation reflects functional importance, CRISPR-mediated disruption of these conserved regions would be expected to yield stronger phenotypes, particularly negative phenotype scores indicative of growth defect, given MYC’s role in promoting cell growth. Paradoxically, we observed the opposite: noncoding base-pairs with significant phenotypes tend to be less conserved than noncoding base-pairs without significant phenotypes (Fig. 3A, P = 3.0x10^-6^), regardless of whether conservation was evaluated using genomes of 100 vertebrates or 30 mammals (mostly primates), measured as phastCons or phyloP scores (Fig. S5A). Across the entire locus, conserved noncoding base-pairs were associated with significantly weaker, rather than stronger phenotypes (Fig. 3B, P = 2.0x10^-20^, similar results using *phyloP* and different conservation threshold in Fig. S5B). This inverse relationship was most pronounced in the promoter region (Fig. 3C), followed by sequences upstream of the promoter and within the downstream 5’ UTR (Fig. S5C), all enriching for transcriptional cis-regulatory elements (Fig. 2D). This is consistent with other studies showing rapid divergence of transcription factor binding sites between species^38^ and that most cis-regulatory elements of transcription lack sequence conservation^39^.

**Figure 3.**
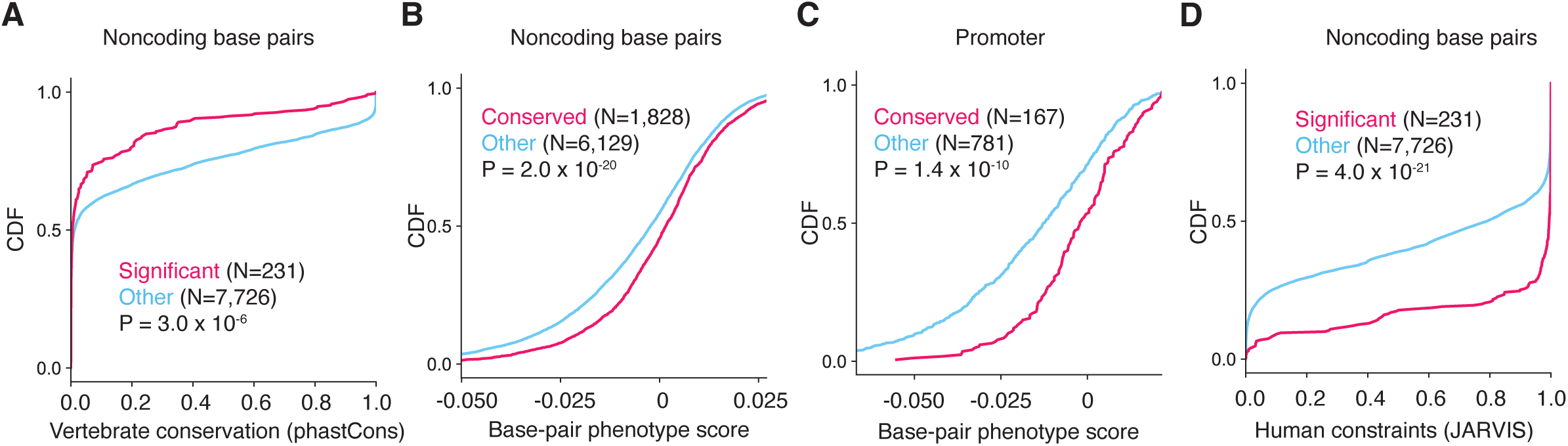
Functional noncoding sequences in *MYC* have reduced vertebrate conservation but increased human-specific constraints. **A**. CDF of conservation scores (phastCons) in 100 vertebrate species comparing significant and other noncoding base-pairs. **B**. CDF of phenotype scores comparing conserved (phastCons >= 0.5) and non-conserved (phastCons < 0.5) noncoding base-pairs. **C**. As in B for base-pairs in the promoter. **D**. As in A for human lineage-specific constraint scores (JARVIS). Kolmogorov–Smirnov test was used to compute all the P values.

The paradox highlights the limitation of relying on evolutionary sequence conservation to identify functional elements in the noncoding genome. Recently, evolutionary sequence conservation agnostic, human lineage-specific variation intolerance scores (e.g., JARVIS^40^) have been developed as an alternative. Indeed, we found that base-pairs with a significant phenotype are under markedly stronger constraint in the human lineage (Fig. 3D). However, overall, JARVIS score only weakly correlates with our phenotype score (*r* = -0.18, Fig. S5D), underscoring the importance of unbiased experimental quantification of phenotypes for noncoding sequences.

Additional evidence suggests that a combination of rapidly evolving 5’ activating DNA elements and deeply conserved 3’ repressive RNA elements may represent a generalizable functional architecture of genes involved in cell growth. Reanalysis of a saturation mutagenesis screen targeting the promoter and 5’ UTR of the ribosomal protein gene *RPS19* in human K562 cells^41^ similarly revealed reduced sequence conservation at base-pairs associated with strong phenotypic effects (Fig. S5E). We also observed a significant negative correlation between 3’ UTR conservation and mRNA half-lives across all human genes (Fig. S5F), extending earlier observations that were limited to a few dozen mRNAs^42,43^. This finding highlights conserved 3’ UTRs as broadly acting post-transcriptional repressive elements.

### An ultraconserved 3’ UTR element indispensable for MYC-addicted cells

Within the largely repressive core 3’ UTR, we identified a single, unique site essential for cell growth (highlighted with a red star in Fig. 1F). Among the sgRNAs targeting this site, sg723 produced the strongest phenotype, which has a GGT PAM and would have been missed by other Cas9 with more restrictive PAM (Fig. 4A). To validate the growth phenotype, we performed a competitive growth assay comparing cell lines stably expressing sg723 or a non-targeting control (sgCtrl). Consistent with the screen results, sg723-expressing cells were progressively and substantially depleted over 10 days when co-cultured with control cells (Fig. 4B).

**Figure 4.**
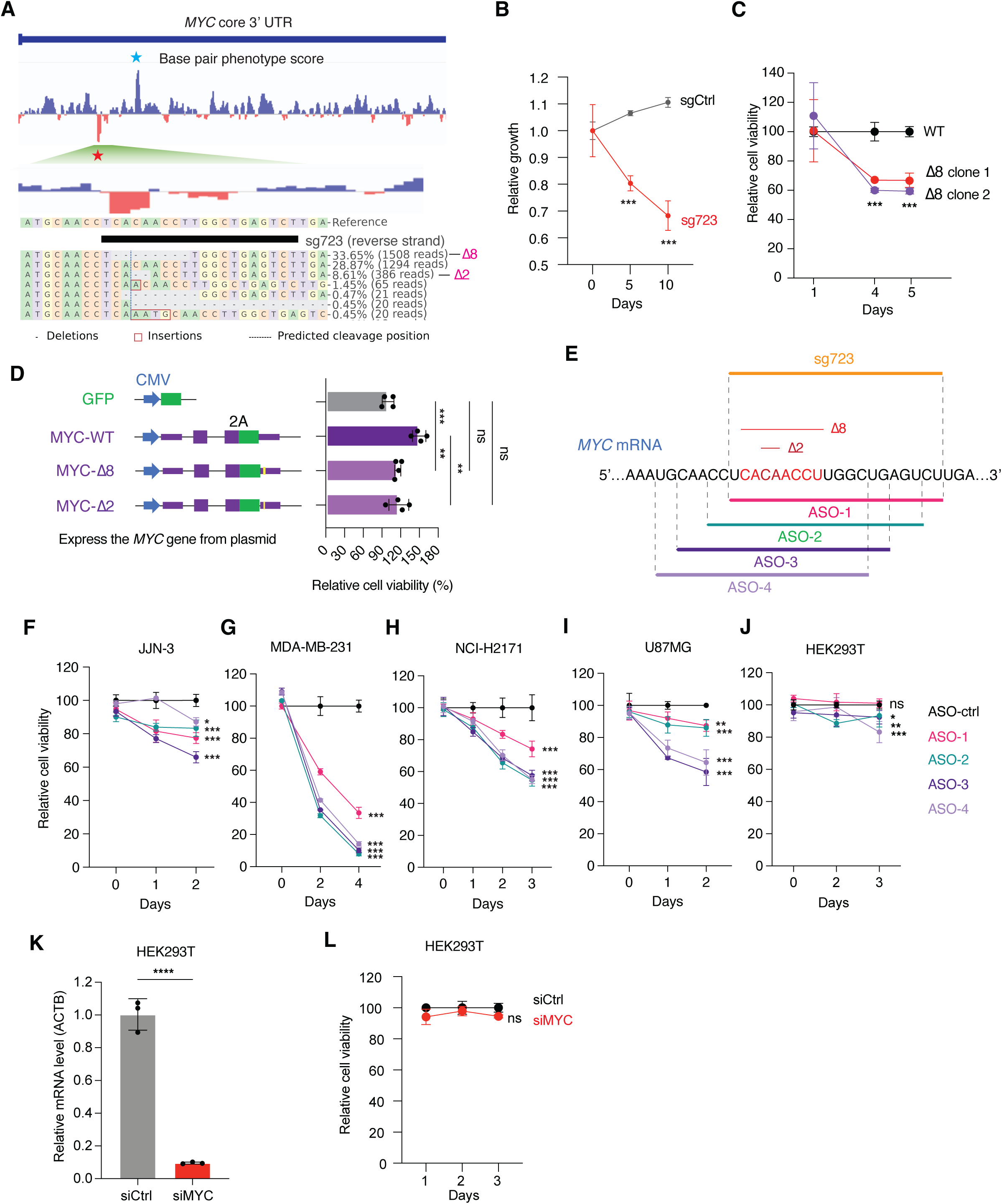
An ultraconserved 3’ UTR element required for MYC-dependent cell growth. **A**. Base-pair phenotype scores of the 3’ UTR and zoom-in to the sg723 target site. Blue and red stars highlight 3’ UTR base pairs with the strongest positive and negative phenotype scores, respectively. Amplicon sequencing reads indicating indels were also shown (only those with frequency > 0.4%). **B**. Competitive growth assay comparing HF-SpRY-Cas9 D11 cells expressing sg723 or a non-targeting control sgRNA. Data were normalized to Day 0. **C**. Relative cell viability (WST-1) comparing two clonal cell lines deleting the ultraconserved element to a cell line targeted by a control sgRNA. Data were normalized to WT. **D**. Relative cell viability (WST-1) comparing cells overexpressing GFP alone, GFP fused to a full-length MYC gene (MYC-WT, MYC-𝛥8, or MYC-𝛥2) using plasmids. Data were normalized to GFP vector. **E**. Four ASOs targeting the ultraconserved element. **F**-**J**. Relative cell viability (WST-1) in indicated cell lines transfected with a control ASO or one of four ASOs targeting the ultraconserved RNA element. Data were normalized to ASO-ctrl. **K**. *MYC* mRNA knockdown with siRNAs in HEK293T cells. Data were normalized to siCtrl. **L**. *MYC* knockdown did not significantly reduce HEK293T cell viability. Data were normalized to siCtrl. ns, not significant, *: p<0.05, **: p<0.01, ***: p<0.001, ****: p<0.0001.

To evaluate potential off-target effects of sg723, we used CasOFFinder to predict candidate off-target sites genome-wide^44^. Among the predicted sites, two contain a single mismatch and thirteen contain two mismatches (Supplementary Table 4). None overlaps genes classified as essential in DepMap^26^, with the exception of one two-mismatch site within *PIK3R5*, which is essential in only ∼1% of CRISPR screens. Targeted amplicon sequencing of top twelve predicted off-target sites, including *PIK3R5,* in SpRY-Cas9/sg723-expressing cells revealed no detectable mutations (Fig. S6). Together, these results indicate that SpRY-Cas9/sg723 exhibits minimal off-target activity and that the observed proliferation defect is attributable to disruption of the targeted region within the *MYC* 3’ UTR.

Sequencing of the target region in sg723-expressing cells revealed a predominant 8-nt CACAACCT deletion (Δ8) and a less frequent 2-nt CA deletion (Δ2) nested within the same region; with the expected cleavage site located after the first CA dinucleotide (Fig. 4A). The same mutational spectrum was observed in both the wild-type (Fig. 4A) and GFP-tagged alleles in the heterozygous D11 cells (Fig. S7A). These mutational patterns were further recapitulated in the triple-negative breast cancer (TNBC) cell line MDA-MB-231 expressing wild-type *MYC* (Fig. S7B). Remarkably, the deleted sequences were unusually conserved: the first 7-nt CACAACC is perfectly conserved in at least 82 vertebrate genomes (Fig. S4A). The let-7/miR-34 target site described earlier is the only other sequence with this level of conservation in the 3’ UTR. Intriguingly, these two ultraconserved sites are only 40-nt apart (red and blue stars in Fig. 1F; see enlarged view in Fig. S4), with the intervening sequence forming a highly conserved stem-loop structure with numerous covariations indicating strong evolutionary pressure in maintaining the RNA structure^45^. The tight clustering of three highly conserved sequence and structural elements with strong but opposing effects suggests possible combinatorial regulation.

We further derived two clonal cell lines carrying the 8-nt CACAACCT deletion (𝛥8) in the *MYC* locus and observed significant growth defect compared to control (Fig. 4C). To further rule out Cas9 off-target effect as a confounding factor of the growth phenotype, we directly tested the growth-promoting role of the 3’ UTR element by transfecting in HEK293T cells plasmids encoding the full-length *MYC* gene with and without the ultraconserved element (Fig. 4D). Briefly, we cloned the entire *MYC* gene in an 8.6-kb genomic DNA into a plasmid, including introns and UTRs (Fig. 4D and Fig. S2B-C). RNA-seq confirmed the expression of a *MYC* transcript from the plasmid with splicing and polyadenylation patterns and efficiency similar to the endogenous transcript (Fig. S2C). Using this wild-type construct (MYC-WT) as a template, we further generated two mutants corresponding to the most frequent outcomes of the sg723 targeting: an 8-nt deletion (MYC-𝛥8) and a 2-nt deletion (MYC-𝛥2) in the 3’ UTR element (Fig. 4D). When transfected in HEK293T cells, only MYC-WT but not MYC-𝛥8 or MYC-𝛥2 significantly promoted cell growth (Fig. 4D).

Together, these convergent observations from bulk-mutated populations, clonal cell lines, and plasmid-expressed mutants validate the screen results and demonstrate that a highly conserved element within the *MYC* 3′ UTR is functionally required to promote cell proliferation.

### 3’ UTR-targeting ASOs inhibit MYC-dependent cancer cells

We hypothesize that the growth-promoting 3’ UTR element could be inhibited by steric-blocking antisense oligos (ASOs) to suppress cell proliferation. To test this, we designed four partially overlapping ASOs targeting this element, including ASO-1, which binds the same RNA region targeted by sg723 on the DNA (Fig. 4E, Supplementary Table 5). All ASOs were fully modified with 2′-O-methoxyethyl (2′-MOE) to enhance stability and binding affinity while reducing cytotoxicity and preventing RNase H-mediated target RNA cleavage. Compared with a negative control ASO (ASO-ctrl), all four ASOs significantly reduced cell viability in the JJN-3 cell line (parental of D11 expressing wild-type MYC without a C-terminal GFP) (Fig. 4F). Similar effects were observed in multiple cancer cell lines that depend on MYC for proliferation (Fig. S8), including MDA-MB-231 (TNBC, Fig. 4G), NCI-H2171 (small cell lung cancer, Fig. 4H), and U87MG (glioblastoma, Fig. 4I), supporting the broader applicability of these ASOs across diverse MYC-dependent cancer types. In contrast, the growth inhibitory effects were markedly attenuated in HEK293T cells (Fig. 4J), a non-malignant human cell line whose proliferation is independent of MYC, as efficient *MYC* knockdown by siRNA (Fig. 4K) does not affect HEK293T cell viability (Fig. 4L). The lack of growth defect in HEK293T cells upon ASO treatment also indicates that the observed phenotypes in other cells (Fig. 4F-I) are unlikely to result from ASO off-target effects or nonspecific toxicity.

Together, these results reveal that the conserved 3’ UTR element is a vulnerability of MYC-dependent cancer cells that can be selectively targeted by ASOs.

### Controlling MYC activity without affecting abundance

Unexpectedly, despite the pronounced growth inhibition observed upon targeting the 3’ UTR element, neither *MYC* mRNA (Fig. 5A) nor protein (Fig. 5B) abundance was significantly altered when cells were targeted with SpRY-Cas9/sg723 in D11 cells. These findings was further supported by the lack of change in GFP levels, which is translated from the same *MYC* mRNA in these cells^24^ (Fig. S9A). Similar results were observed in MDA-MB-231 cells (Fig. S9B), further confirmed using five different commercially available MYC antibodies (Fig. S9C). The lack of significant change in MYC protein abundance was also observed in MYC-𝛥8 clonal cells (Fig. S9D).

**Figure 5.**
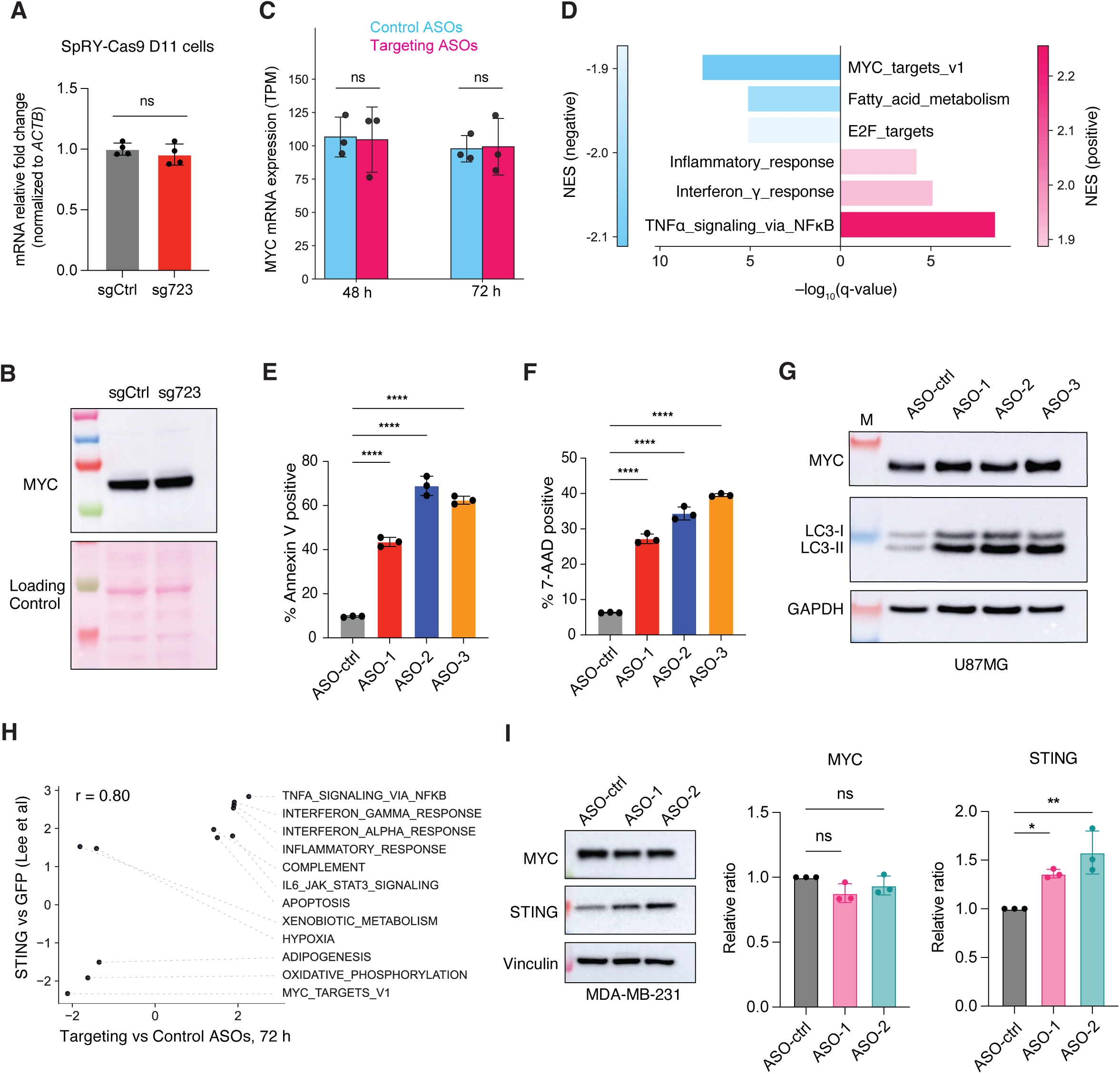
The ultraconserved 3’ UTR element controls MYC activity without affecting its abundance. **A**. RT-qPCR quantification of *MYC* mRNA in cells targeted by sg723 or a non-targeting control (sgCtrl) as in Fig. 4B. Data were normalized to sgCtrl. **B**. As in A but for MYC protein. Ponceau S was used as loading control. **C**. *MYC* expression measured by RNA-Seq in TPM (transcripts per million) 48 or 72 h after transfection with three targeting ASOs and three control ASOs. Significance reflects adjusted p-values from DESeq2 differential expression analysis performed on raw counts. **D**. Top three MSigDB hallmark gene sets significantly enriched in up- or down-regulated genes (ASO treated vs ASO control, 72 h). NES: normalized enrichment scores. **E**-**F**. Flow cytometric analysis of apoptosis in MDA-MB-231 cells following ASO treatment. Cells were collected 72 h post-transfection and stained with Annexin V (E) and 7-AAD (F). **G**. Western blotting of MYC and LC3 48h after ASO treatment. M: Marker. **H**. Scatter plot of normalized enrichment scores (NES) of MSigDB hallmark gene sets significant (p.adjust < 0.05) in this study (x-axis) and Lee *et al*. **I**. Western blotting for MYC and STING 72h after ASO treatment. Data were normalized to ASO-ctrl. ns, not significant, *: p<0.05, **: p<0.01, ****: p<0.0001.

To elucidate how this 3’ UTR element regulates cell growth independently of MYC abundance, we profiled transcriptomic changes in MDA-MB-231 cells transfected with three ASOs targeting the 3’ UTR element (ASO-1, ASO-2, and ASO-3, Fig. 4E) compared with three distinct control ASOs (Supplementary Table 5). Differential gene expression analysis at 48 h (Supplementary Table 6) and 72 h (Supplementary Table 7) using DESeq2^46^ confirmed that *MYC* expression remained unchanged by ASO treatment (Fig. 5C). Remarkably, Gene Set Enrichment Analysis (GSEA) identified MYC target genes as the most down-regulated MSigDB hallmark gene set^47^ (Fig. 5D). This finding was corroborated by independent analyses separating MYC-upregulated and MYC-downregulated gene sets^48^: MYC-upregulated genes were decreased, whereas MYC-downregulated genes were increased, in cells treated with ASOs targeting the 3’ UTR element (Fig. S10A). These results indicate that steric blocking of the 3’ UTR element by ASOs inhibited MYC activity as a transcription factor despite unaltered MYC expression.

Consistent with the observed growth defects upon inhibition of the 3’ UTR element, hallmark gene sets associated with cell cycle progression and metabolism were strongly down-regulated, while apoptosis was significantly upregulated (Fig. S10B). Indeed, Annexin V (Fig. 5E) and 7-AAD (Fig. 5F) staining confirmed that ASO treatments robustly induced apoptosis. We also observed a significant increase in autophagy (Fig. 5G).

Intriguingly, the most up-regulated hallmark gene sets were dominated by interferon and innate immune responses (Fig. 5D and Fig. S10B), suggesting a possible mechanism driving cellular apoptosis and autophagy (Fig. 5E-G). Remarkably, these pathway activity changes closely mirrored those observed upon expressing STING (stimulator of interferon genes) in another TNBC cell line MDA-MB-436 (*r* = 0.8, Fig. 5H)^49^. MYC has been reported to suppress innate immune response by directly inhibiting the expression of STING^50^, a key regulator of the cellular innate immune response. In TNBC MDA-MB-436 and SUM159PT Taxol-resistant cells, this is through binding and silencing an intronic enhancer of STING^49^. Consistent with this model, we confirmed a significant increase of STING protein (Fig. 5I) and mRNA (Fig. S10C) in cells treated ASOs targeting the 3’ UTR element. MYC also suppresses interferon-dependent genes by inhibiting TANK-binding kinase 1 (TBK1)^51,52^, which was also up-regulated in ASO-treated cells (Fig. S10D).

Together, these results indicate that the 3’ UTR element is essential for maintaining MYC’s function as a transcription factor, particularly its repression of innate immune genes. Targeting this element with ASOs functionally inhibits MYC activity and consequently induces interferon response, resulting in apoptosis, autophagy, and cell growth defects, without affecting *MYC* mRNA and protein expression.

### The 3’ UTR element regulates *MYC* mRNA and protein localization

To elucidate how the 3’ UTR element regulates MYC function without altering its protein abundance, we examined MYC protein subcellular localization by immunofluorescence in MDA-MB-231 cells. In cells transfected with control ASO, MYC protein localized predominately to the nucleus (Fig. 6A), consistent with its role as a transcription factor. In contrast, transfection of three ASOs targeting the 3’ UTR element consistently caused a marked redistribution of MYC to the cytoplasm, accompanied by a reduction in nuclear MYC in many cells (Fig. 6A, quantified in Fig. 6B). Given that MYC functions as a transcription factor in the nucleus, this mislocalization likely underlies the observed impairment of MYC function and associated growth defects, particularly in MYC-addicted cancer cells.

**Figure 6.**
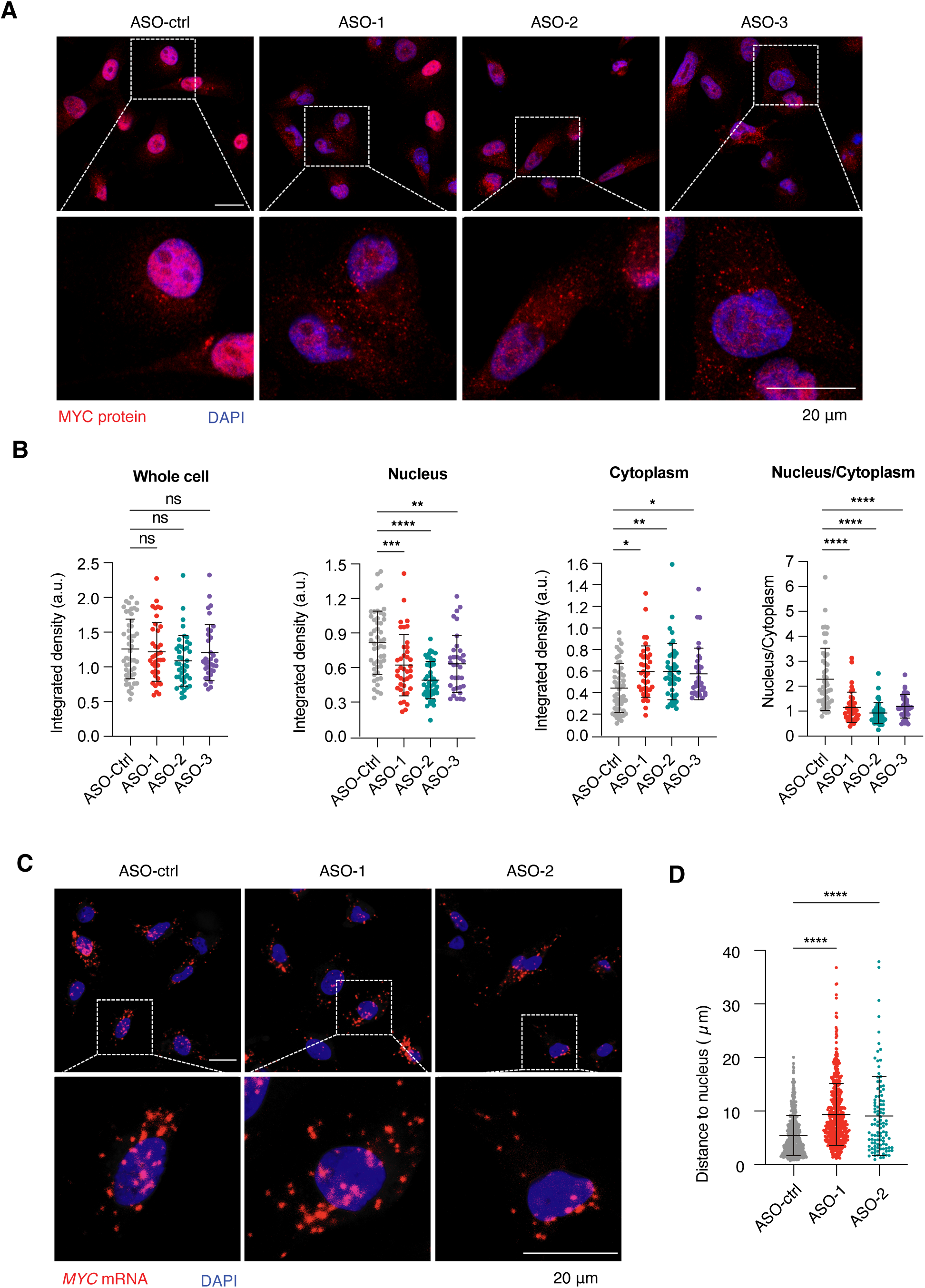
The ultraconserved 3’ UTR element facilitates *MYC* mRNA and protein localization. **A**. Representative immunofluorescence images of MYC protein in MDA-MB-231 cells 48 h after transfection with ASO-ctrl or three ASOs targeting the 3’ UTR element. **B**. Quantitative analysis of fluorescence shown in A. a.u.: Arbitrary units. ASO-ctrl: N=49; ASO-1: N=39; ASO-2: N=42; ASO-3: N=34. **C**. Representative RNAScope images showing *MYC* mRNA localization in MDA-MB-231 cells. **D**. Quantification of C with the distance of each *MYC* mRNA spot to the nearest nucleus. ASO-ctrl, N=483; ASO-1, N=500; ASO-2, N=104. ns, not significant, *: p<0.05, **: p<0.01, ***: p<0.001, ****: p<0.0001.

To directly test the role of the 3’ UTR element in MYC protein localization, we inserted an N-terminal 2xHA tag to MYC expressed from plasmids (as in Fig. 4D) to distinguish it from endogenous MYC protein. Upon transient transfection into U87MG cells, immunofluorescence for the HA tag revealed that while MYC-WT localized primarily to the nucleus, both MYC-𝛥8 and MYC-𝛥2 mutants exhibited pronounced cytoplasmic accumulation in many cells (Fig. S11). These results demonstrate that the 3’ UTR element promotes nuclear localization of MYC protein.

We next investigated the mechanism by which the 3’ UTR element facilitates MYC protein nuclear localization. Previous studies have shown that *MYC* mRNA localizes near the nuclear periphery, where local translation facilitates efficient nuclear import of the nascent MYC protein, a process particularly important given MYC protein’s short half-life (20-30 minutes)^53^. Using RNAScope to visualize endogenous *MYC* mRNA, we confirmed predominant perinuclear localization of *MYC* mRNA in control ASO-treated cells (Fig. 6C). In contrast, many cells transfected with two different MYC ASOs displayed a dispersed cytoplasmic distribution of *MYC* mRNA, with increased distance from the nucleus (Fig. 6C). Compared with MYC proteins synthesized from perinuclear mRNAs, those translated from distal mRNAs are more susceptible to degradation prior to nuclear import. This mislocalization of *MYC* mRNA likely accounts for reduced nuclear MYC protein (Fig. 6A-B) and impaired its normal function in the nucleus.

## DISCUSSION

In this study, we performed, to our knowledge, the first saturated, base-pair-resolution functional dissection of a complete endogenous gene in its native genomic context. By mapping the fitness landscape of the *MYC* locus, we delineated the relative contributions of coding and noncoding sequences to gene function, revealed the regulatory logic of the locus, and uncovered a novel RNA regulatory element that can be targeted to inhibit cancer cells.

Central to these discoveries is the ability of the high-fidelity, near-PAMless SpRY-Cas9 to target virtually every base-pair with multiple sgRNAs (Fig. 1B). This enables an unbiased, saturating, base-pair resolution dissection of genomic function while minimizing the influence of individual sgRNA off-target effects and variable on-target efficiencies by aggregating signals from multiple sgRNAs targeting the same base-pair (Fig. 1B). The power of the SpRY-Cas9 system is evident from the observation that most of the top ranked sgRNAs (86.3%) do not target canonical NGG PAM sites (Fig. 1I), including sg723, which led to the identification of the druggable ultraconserved 3’ UTR element (Fig. 4-6). Together, these results establish high-fidelity SpRY-Cas9 as a robust platform for saturation mutagenesis, expand the functional genomics toolkit for mapping gene regulatory architecture with unprecedented resolution, and accelerate therapeutic discovery from the noncoding genome.

Our unbiased analysis reveals the critical role of noncoding sequences and the functional architecture of the *MYC* locus. We find that noncoding regions account for approximately two-thirds of the base pairs with the strongest phenotypic effects (Fig. 2A). Because this analysis was performed in a single cell line and under unstimulated conditions, this proportion is likely an underestimate, as many noncoding regulatory elements act in a cell type- or condition-specific manner. These noncoding regulatory elements segregate into two distinct domains: a rapidly evolving 5′ region that activates *MYC* transcription, and a highly conserved 3′ region that represses *MYC* post-transcriptionally (Fig. 2D). The opposing activities of these two domains provide a mechanistic explanation for the paradoxical anticorrelation between evolutionary conservation and phenotypic impact at this locus (Fig. 3). Extending this approach to additional genes will be essential to determine the generality of these principles.

Another key finding from our screen was the identification of an ultraconserved 3’ UTR element essential in MYC-dependent cancer cells (Fig. 4-5). Disruption of this region did not alter *MYC* mRNA or MYC protein levels but instead mislocalized *MYC* mRNA from the nuclear periphery to the cytoplasm, resulting in MYC protein accumulation outside the nucleus and diminished transcriptional activation of MYC target genes (Fig. 6). In addition to the decrease in nuclear abundance of MYC, recent studies demonstrated that disrupting *MYC* mRNA perinuclear localization and translation in the TIS granule^54^ also inhibits MYC activity by disrupting co-translational protein-protein interactions^55^. Together, these findings highlight subcellular RNA trafficking and localized translation as an important and underappreciated layer of gene regulation, particularly for short-lived, nuclear proteins such as MYC. Future studies will seek to identify protein factors or other regulatory RNAs that recognize the 3’ UTR element and mediate the perinuclear localization of the *MYC* mRNA, as well as to explore potential combinatory regulations involving the adjacent ultraconserved dual miRNA target site with opposing effects (Fig. S4), which is separated by a deeply conserved RNA structure^45^.

The maturation of ASO technology into a robust, programmable drug discovery platform has enabled the development of a broad portfolio of approved therapies, transforming the treatment landscape for both rare and common diseases through precise modulation of RNA function^56,57^. We developed steric blocking ASOs that target the ultraconserved *MYC* 3′ UTR element and selectively eliminate MYC-dependent cancer cells (Fig. 4). Previously developed *MYC*-targeting ASOs – such as INX-3280, INX-6295^58^, AVI-4126^59^, and MYCASO^60^, act via RNase H–mediated cleavage of *MYC* RNA, which are associated with widespread off-target RNA degradation and immunogenicity^61,62^. Similar limitations apply to *MYC* RNA knockdown using siRNAs. In contrast, fully 2′-MOE-modified steric blocking ASOs, as used in this study, exhibit substantially reduced immunogenicity^63^, and by acting through steric hindrance rather than RNA cleavage, avoid off-target RNA degradation^64^. The minimal to moderate growth effects observed in MYC-independent, non-malignant HEK293T cells following ASO treatment (Fig. 4J–L) indicate that these ASOs do not elicit significant off-target effects or nonspecific cytotoxicity. While these ASOs may still bind unintended RNAs, such interactions are generally less deleterious than cleavage-mediated degradation. Compared to protein-targeting drugs like OMO-103^65^, RNA-targeted ASOs may offer improved intracellular delivery, as they do not require nuclear entry. Beyond its well-established role in cancer, MYC is a critical player in many diseases. Thus, our ASOs may also have therapeutic potential in MYC-related conditions such as metabolic disorders^65^, obesity^66^, cardiovascular diseases^67–69^, autoimmune diseases^70,71^, neurological disorders^72^, and aging^73^.

Our work establishes a blueprint for unbiased, high-resolution interrogation of the genome. We anticipate that extending this approach to additional genes will collectively reveal fundamental design principles of gene structure and regulatory architecture, while uncovering previously unrecognized layers of gene regulation. Our study also highlights the power of coupling unbiased functional screens with RNA-targeted modalities such as ASOs to accelerate translational applications, including the development of therapies against targets long considered “undruggable” by conventional approaches.

## ACKNOWLEDGMENTS

We thank members of the Cardiometabolic Genomics Program, the Chaolin Zhang group, and the Sara Zaccara group at Columbia University for discussion. We thank Dr. Jerry Pelletier and Dr. Francis Robert for sharing the D11 and JJN-3 cells, and Dr. Peter S. Sims for sharing the U87MG cells. X.W. is supported by NIH Director’s New Innovator Award (1DP2GM140977), Pershing Square Sohn Prize for Cancer Research, and Pew-Stewart Scholar for Cancer Research Award. This research was funded in part through the NIH/NCI Cancer Center Support Grant P30CA013696 and used the Genomics and High Throughput Screening Shared Resource, the Molecular Pathology Shared Resource, and CCTI Flow Cytometry Core. The CCTI Flow Cytometry Core is supported in part by the Office of the Director, National Institutes of Health under awards S10OD030282 and S10OD020056.

## AUTHOR CONTRIBUTIONS

P.S. and X.W. conceived the project. P.S. performed most of the experiments. F.Y. performed most of the computational analyses. X.W. supervised the project. Tala assisted with RT-qPCR, flow cytometry, immunofluorescence, RNAScope and plasmids generation, supervised by J.Q. W.H. assisted in confirming ASO function. A.O.A. assisted with plasmids generation and validation as well as confirmation of ASO function. C.H.K. assisted with computational analyses. P.S., F.Y., X.W. wrote the manuscript with input from all authors. A.T., M.R.M., M.P.R., E.R.R., and L.X. edited the manuscript and provided critical feedback.

## METHODS

### Cell culture

HEK293T, JJN-3, D11, MDA-MB-231, U87MG, NCI-H2171 cell lines were used in this research. HEK293T, MDA-MB-231 and NCI-H2171 cells were purchased from ATCC. JJN-3 and D11 cells were gifts from Jerry Pelletier. U87MG cell line was gift from Peter A. Sims. HEK293T, MDA-MB-231, and U87MG cells were cultured in DMEM with 4.5 g/L D-glucose (Gibco, 11995073) supplemented with 10% heat inactivated fetal bovine serum (FBS, Gibco, 10438026), no antibiotic was added. JJN-3, D11, and NCI-H2171 cells were cultured in RPMI 1640 Medium (Gibco, 11875119) supplemented with 10% heat inactivated FBS, no antibiotic was added. Low passage number cells were used and cells were passaged upon reaching 80–90% confluency. Cells were maintained at 5% CO2 and 37 °C. All cells used in this study were confirmed to be negative for mycoplasma contamination and routinely tested using the MycoAlert Mycoplasma Detection Kit (Lonza, LT07-418).

### Plasmids construction

The LentiV-HF1-SpRY-Cas9 WT plasmid was generated through a multi-step cloning strategy. First, the LentiV_ABE8e-HF1-SpRY_puro plasmid was constructed by assembling fragments derived from LentiV_Cas9_puro (Addgene, 108100) and pCMV-SpRY-HF1-ABE8e (Addgene, 185670) using the Gibson Assembly® Master Mix (NEB, E2611S) according to the manufacturer’s instructions. The resulting construct contained the HF1-SpRY-ABE8e cassette under the lentiviral backbone control. To generate the LentiV-HF1-SpRY-Cas9 WT plasmid, site-directed mutagenesis was performed to remove the ABE8e domain and revert the D10A nickase mutation in Cas9 to the wild-type catalytic residue. Mutagenic primers were designed using NEBaseChanger (New England Biolabs), and PCR amplification was initially carried out using the Phusion® High-Fidelity DNA Polymerase Kit (NEB, E0553). The resulting amplicons were then processed using the Q5® Site-Directed Mutagenesis Kit (NEB, E0554S) according to the manufacturer’s instructions to introduce the desired mutations and generate the final construct. For sgRNA cloning, we used the lentiU6-sgRNA/EF1a-mCherry (Addgene, 114199), which contains BbsI sites for spacer insertion. For each target, two complementary oligonucleotides were synthesized: a forward oligo (5′-CACC–GN_20_–3′) identical to the target RNA sequence and a reverse oligo (5′-AAAC–reverse complement of N_20_C–3′). The vector was digested with BbsI-HF (NEB, R3539S) and the correct band was purified using the Macherey-Nagel™ Gel and PCR Clean-up Kit (Macherey-Nagel, 740609). Oligo pairs were annealed (95°C for 5 min, ramped to 25°C at 5°C/min), and ligated into the digested backbone using T4 DNA ligase (NEB, M0202T). For plasmid transformation, all plasmids were transformed into NEB Stable Competent E. coli (C3040H) cells via heat shock at 42°C for 30 seconds. Cells were recovered in NEB Outgrowth Medium for 1 h at 30°C with shaking (250 rpm) and plated onto selective LB-agar plates containing the appropriate antibiotic. Plates were incubated at 30°C overnight to minimize potential recombination events in lentiviral backbones. Individual colonies were picked and cultured in 5 mL of LB broth containing the corresponding antibiotic at 30°C for 24 h with shaking. Plasmid DNA was purified using the Macherey-Nagel™ Nucleospin™ Plasmid Transfection-grade Kit (Macherey-Nagel, 740490) according to the manufacturer’s protocol. Final construct verification was conducted through full-length plasmid sequencing performed by Plasmidsaurus to confirm each plasmid sequence.

### Cloning the *MYC* locus on plasmids

A full-length *MYC* expression construct was generated to include the endogenous 5′ untranslated region (5′UTR), all introns and exons, with or without a 2A-GFP cassette, the complete 3′UTR, and an additional 150 bp downstream of the 3′UTR in a pCMV-based backbone. Plasmids were assembled using a seamless Gibson-style cloning strategy. The pCMV-GFP backbone (Addgene, 11153) and the full-length *MYC* genomic fragment were amplified by PCR using Platinum™ SuperFi™ II DNA Polymerase (Invitrogen, 12361010). During vector amplification, the GFP sequence in the pCMV-GFP backbone was excluded. Primers were designed using the GeneArt Seamless Cloning and Assembly online design tool and are listed in Supplementary Table 2. PCR reactions (50 µL total volume) contained 1× SuperFi™ II Buffer, 0.2 µM each primer, 200 µM each dNTP, Platinum™ SuperFi™ II DNA Polymerase, template DNA, and nuclease-free water to volume. Templates included 20 ng pCMV-GFP plasmid DNA for vector amplification and 200 ng D11 genomic DNA for *MYC* amplification. PCR was performed for 30 cycles with extension times of 4 min for the vector backbone and 9 min for the full-length *MYC* insert. PCR products were resolved by agarose gel electrophoresis, and bands of the expected size were excised and purified using the Macherey-Nagel™ Gel and PCR Clean-up Kit (Macherey-Nagel, 740609) according to the manufacturer’s instructions. Purified PCR products were assembled using the GeneArt® Seamless Cloning and Assembly Enzyme Mix (Invitrogen, A14606). Assembly reactions were set up at a molar ratio of *MYC* insert to vector of 2:1. Insert DNA and linearized vector were combined with nuclease-free water to a final volume of 5 µL, followed by the addition of 5 µL GeneArt 2× Enzyme Mix. Reactions were mixed gently, briefly centrifuged, and incubated at room temperature for 15–30 min. Following incubation, the same methods for bacterial transformation, plasmid purification, and full-length plasmid sequencing as described above were used.

For generation of the pCMV-MYC-2A-d2GFP-Δ8 and pCMV-MYC-2A-d2GFP-Δ2 plasmids, pCMV-MYC-2A-d2GFP was used as the parental backbone. Gibson-style seamless cloning was performed following the same procedures described above, using primers listed in Supplementary Table 2.

For generation of the pCMV-MYC-2A-d2GFP-ΔAlu and pCMV-MYC-ΔAlu plasmids, pCMV-MYC-2A-d2GFP and pCMV-MYC were used as the parental backbone. PCR primers were designed using NEBaseChanger (New England Biolabs), and all primer sequences are listed in Supplementary Table 2. Site-directed mutagenesis was performed by PCR using Platinum™ SuperFi™ II DNA Polymerase (Invitrogen, 12361010) with 10 ng of pCMV–Whole MYC plasmid as template in a 50 µL reaction containing 1× SuperFi™ II Buffer, 200 µM each dNTP, and 0.2 µM of each primer. PCR was performed for 30 cycles with a 12 min extension step. Following amplification, PCR products were subjected to kinase–ligase–DpnI (KLD) treatment using the Q5® Site-Directed Mutagenesis Kit (New England Biolabs, E0554S) by combining 1 µL PCR product with 5 µL 2× KLD Reaction Buffer, 1 µL 10× KLD Enzyme Mix, and 3 µL nuclease-free water, followed by incubation at room temperature for 5 min. Following incubation, the same methods for bacterial transformation, plasmid purification, and full-length plasmid sequencing as described above were used.

All oligonucleotides were synthesized by Integrated DNA Technologies (IDT).

### sgRNA library design

The 10,055-bp genomic locus (chr8:127,734,053-127,744,107, hg38) containing the *MYC* gene was used for saturation mutagenesis. This region covers all annotated *MYC* transcript isoforms (GENCODE v49) and over 1-kb intergenic DNA on each end containing evolutionary conserved sequences across 100 vertebrate genomes (Fig. 1A). All possible 20-nt sequences on both the plus and minus strands in this region were used as sgRNA spacers. In addition, genomic DNA amplicon sequencing identified three SNPs in the D11 cells used. All 20-nt sequences overlapping with these SNPs on either strand were also added to the library. In the D11 cell line, one of the *MYC* alleles contains P2A-GFP at the C-terminal end (before the stop codon). All 20-nt sequences overlapping with this insert (both plus and minus strands) were also included in the library. We also included 565 negative control sgRNAs, including 404 targeting a 221-nt random sequence, 111 targeting the non-essential AAVS1 locus, and 50 non-targeting sgRNAs. In total, 22,623 sgRNAs was included for this project.

### sgRNA library cloning

A DNA oligo pool containing all 20-nt sgRNA spacer sequences and constant flanking sequences (5’-TAACTTGCTATTTCTAGCTCTAAAAC and CGGTGTTTCGTCCTTTCCACAAGATATATAAAGCCA-3’) was synthesized by Twist Bioscience. The oligo pool was resuspended in TE buffer (10 mM Tris-Cl pH 8.0, 1 mM EDTA) to 10 ng/ul and PCR-amplified (10 ng template, 10 cycles) using the KAPA HiFi HotStart PCR Kit (Kapa Biosystems, KK2502) and primers listed in Supplementary Table 2. The PCR product was cleanup using the Macherey-Nagel™ Gel and PCR Clean-up Kit (Macherey-Nagel, 740609). NEBuilder® HiFi DNA Assembly Master Mix was used to insert the sgRNA library into the sgRNA vector lentiU6-sgRNA/EF1a-mCherry (Addgene, 114199) via Gibson assembly. The product is cleaned up using isopropanol precipitation to remove salts that can interfere with electroporation. A total of 550 ng DNA was used in electroporation in Endura competent cells (Endura, 60242-1). Bacteria were plated in six 245 mm square dishes and cultured for 18 h. Bacterial colonies were scraped off in liquid LB, followed by maxiprep with MACHEREY-NAGEL kit (Macherey-Nagel, 740414.50).

### CRISPR saturation mutagenesis screening

#### HF1-SpRY-Cas9 stable cell line generation

We generated the HF1-SpRY-Cas9 lentivirus using the packaging system consisting of pCMV-dR8.91 and pMD2.G packaging system (gifts from Jonathan Weissman). Packaging plasmids were co-transfected into HEK293T cells in 10 cm dish using Lipofectamine 3000 (Thermo Fisher Scientific, L3000015) according to the manufacturer’s instructions. The culture medium was replaced with fresh growth medium 6 h post-transfection. Viral supernatants were collected at 48 and 72 h post-transfection, followed by centrifugation at 800 g for 5 min to remove cell debris. The clarified supernatant containing the lentivirus was filtered through a 0.45 μm PVDF membrane filter (Sigma, SLHV033RS), concentrated using Lenti-X™ Concentrator (Takara, 631231), resuspended in RPMI 1640 media, and used to infect 2.1×10^6^ cells in the presence of polybrene (8 ug/ml). Media was changed 24 h after infection. Three days after infection, cells were selected with puromycin (0.5 ug/ml) for 4 days. The expression of HF1-SpRY-Cas9 was confirmed by RT-qPCR at the end of the selection.

#### Lentivirus for sgRNA

sgRNA library lentivirus was prepared using the same method for the HF1-SpRY-Cas9 stable cells generation. 30.1 ug of lentiviral plasmid was used.

#### Infection and screening

For functional interrogation, 5×10⁶ cells were seeded per T75 flask (20 mL per flask), across 50 T75 flasks, yielding a total of 2.5×10⁸ cells for infection. Cells were transduced at a low multiplicity of infection (MOI < 0.3) to ensure that the majority of cells received only a single gRNA cassette. Lentivirus was added at 0.2 mL per 5×10⁶ cells in 20 mL total culture media, supplemented with 16 µL polybrene (8 µg/mL). After 24 h, media were replaced without discarding cells; cells were washed three times with fresh growth media and centrifuged at 250 g for 5 min to collect all cells. At least 2.5×10⁸ cells were maintained continuously for culture. After 48 h, the sgRNA positive ratio was tested by flow cytometry testing the mCherry. Take the day that after 48 h infection as Day 0. At the Day 6, sorted the mCherry positive cells, keep cells about ∼1000× library coverage in culture. Collect cells every other day to Day 20.

For H-MOI infection. 5x10^6^ cells per T75 flask, 6 Flasks, 3×10^7^ cells in total. MOI is 8.7, polybrene (8 µg/mL). After 24 h, media were replaced without discarding cells; cells were washed three times with fresh growth media and centrifuged at 250 g for 5 min to collect all cells. After 48 h, the sgRNA positive ratio was tested by flow cytometry testing the mCherry. Take the day that after 48 h infection as Day 0. Collect all cells for DNA extraction.

#### Genomic DNA Extraction

For day 0, day 8 and day 20 samples, genomic DNA was isolated using the Macherey-Nagel NucleoSpin Tissue kit (Macherey-Nagel, 740952.50) according to the manufacturer’s protocol with modifications. To ensure DNA purity, samples were treated with RNase A to eliminate RNA contamination. DNA was eluted using 50 μL elution buffer (EB) per column with a 3 min incubation at room temperature, followed by centrifugation for 1 min. To maximize DNA recovery, each column was eluted three times, and eluates were pooled. DNA concentration and purity were determined using a NanoDrop spectrophotometer (Thermo Scientific). Sample quality was assessed by measuring absorbance ratios at A_260_/A_280_ and A_260_/A_230_. Purity ratios were: A_260_/A_280_ = 1.86-1.90 and A_260_/A_230_ = 2.41-2.43. For PCR template preparation, DNA concentrations were verified using a Qubit fluorometer (Thermo Scientific) for accurate quantification.

#### PCR Amplification and Purification

Targeted genomic regions were amplified by PCR using extracted genomic DNA as template. PCR reactions were optimized for each sample type with template amounts of 4 μg per 100 μL reaction. Forward primer pools were prepared by combining equal volumes of individual forward primers (100 μM stock concentrations) and diluting the mixture to a final concentration of 10 μM for PCR use. PCR cycling conditions were optimized through preliminary experiments, and the number of amplification cycles was limited to a maximum of 19. Multiple parallel PCR reactions were performed to generate sufficient amplicon for high-throughput sequencing. PCR products from all parallel reactions were pooled and thoroughly mixed. Initial purification was performed using the MN NucleoSpin PCR Clean-up kit (Macherey-Nagel, 740609). Each column was eluted three times with nuclease-free water to maximize product recovery. Purified PCR products underwent gel purification using 1.5% agarose gel electrophoresis in 1× TAE buffer. Target bands were excised and purified using the MN NucleoSpin Gel and PCR Clean-up kit (Macherey-Nagel, 740609). Final purified products were eluted three times per column. Sample quality was assessed by NanoDrop spectrophotometry.

#### Library Preparation and High-Throughput Sequencing

Sequencing libraries were prepared and quantified using the KAPA Library Quantification Kit (Roche, 07960336001) according to the manufacturer’s instructions. Libraries were sequenced on the AVITI sequencing platform (Element Biosciences) using the 2×75 High Output kit. Library normalization and denaturation were performed using IDTE buffer (pH 8.0, Integrated DNA Technologies) as the low TE buffer. Sequencing was performed according to the AVITI System User Guide with paired-end 75 bp reads.

### Competitive assay

To validate the growth-inhibitory effect of individual sgRNAs, we performed a competitive growth assay. We first generated a stable HF-SpRY-Cas9-expressing cell line, then transduced it with either a non-targeting control sgRNA (sgCtrl) or individual targeting sgRNAs to establish separate stable cell populations. Equal numbers of sgCtrl and individual targeting sgRNAs cells (1×10^6^ each) were mixed, and 2×10^5^ cells were collected on the same day as the initial time point (Day 0). The remaining mixed cells were cultured, and 1×10^6^ cells were collected every other day for analysis. Genomic DNA was extracted from each sample, followed by PCR amplification of the sgRNA region and amplicon sequencing (Plasmidsaurus) to quantify the abundance of each sgRNA over time.

### Cell proliferation assays (WST-1)

Cells were incubated with the 10% WST-1 reagent (Sigma,11644807001) for 2 h. The absorbance of the samples was measured using a microplate reader at 440 nm.

### Amplicon sequencing for on-target and off-target mutation analysis

Following Cas9–sgRNA–mediated mutagenesis, genomic DNA was extracted and the target loci were amplified by PCR using Q5 High-Fidelity 2× Master Mix (New England Biolabs, M0492S) with primers listed in Supplementary Table 2. PCR products were resolved by agarose gel electrophoresis and purified prior to sequencing using Plasmidsaurus amplicon sequencing service. Resulting FASTQ files were retrieved and analyzed using CRISPResso2 to quantify insertion–deletion (indel) frequencies and editing outcomes^74,75^. For sg723 off-target analysis, all top 15 predicted off-target sites with less than 3 mismatches were tested except three intergenic sites with two mismatches. These three sites share the same targeting sequence as another site we tested (LINC01145), which showed no mutations.

### ASO transfection

Lyophilized ASOs were centrifuged briefly upon receipt and stored dry at −20°C until use. ASOs were resuspended in nuclease-free water to generate concentrated stock solutions (typically 100–200 µM), calculated based on the manufacturer-provided oligonucleotide amount. Resuspension was performed by gentle pipetting without vortexing, followed by a short incubation at room temperature to ensure complete dissolution. Resuspended ASOs were aliquoted into single-use volumes and stored at −80°C. For experimental use, aliquots were thawed on ice, mixed gently, and diluted to working concentrations. Cells were seeded in 6-well plates with 2 ml media one day prior to transfection. ASOs were transfected at a final concentration of 100 nM. Opti-MEM I Reduced-Serum Medium (Gibco, 31985062) was pre-warmed at 37 °C for 30 min before use. For each well, 7.5 μl of RNAiMAX reagent was diluted in 125 μl of Opti-MEM and mixed with 200 pmol of ASO diluted in another 125 μl of Opti-MEM. Nuclease-free tubes were used for transfection. After incubation for 5 min at room temperature, the ASO-lipid complex was added to the cells. The culture medium was not changed post-transfection. Cells were harvested 48 h or 72 h later for experiments. For WST-1 assays, cells were transfected in 48-well plates using the same procedure and a final ASO concentration of 100 nM. The ASO sequences are listed in Supplementary Table 5.

### siRNA Transfection

HEK293T cells were transfected with either a non-targeting control siRNA pool (Horizon Discovery, D-001810-10-05) or siRNA pools targeting human *MYC* (Horizon Discovery, L-019634-00-0005), using Lipofectamine™ RNAiMAX Transfection Reagent (Invitrogen, 13778) according to the manufacturer’s instructions. Cells were seeded in 6-well plates with 2 ml media one day prior to transfection. siRNAs were transfected at a final concentration of 20 nM. Opti-MEM I Reduced-Serum Medium (Gibco, 31985062) was pre-warmed at 37 °C for 30 min before use. For each well, 7.5 μl of RNAiMAX reagent was diluted in 125 μl of Opti-MEM and mixed with 40 pmol or of siRNA diluted in another 125 μl of Opti-MEM. Nuclease-free tubes were used for transfection. After incubation for 5 min at room temperature, the siRNA-lipid complex was added to the cells. The culture medium was not changed post-transfection. Cells were harvested 48 h later for RNA extraction and RT-qPCR analysis. For WST-1 assays, cells were transfected in 48-well plates using the same procedure and a final siRNA concentration of 20 nM.

### Quantitative reverse transcription PCR (RT-qPCR)

Total RNA was extracted using the MACHEREY-NAGEL NucleoSpin RNA Plus kit (Macherey-Nagel, 740984.250), and cDNA was synthesized using SuperScript™ IV Reverse Transcriptase (Invitrogen, 18090050). Quantitative RT-PCR was performed using SYBR Green Master Mix (Applied Biosystems, A25741) on a QuantStudio 7 Flex Real-Time PCR System (Applied Biosystems). Gene expression levels were normalized to samples transfected with control siRNA or control ASO. ACTB was used as internal control.

### Western Blot

Cells were lysed in ice-cold RIPA buffer (Sigma-Aldrich, R0278) supplemented with protease inhibitor cocktail (Roche, 4693132001). Briefly, culture dishes were placed on ice, cells were harvested and pelleted by centrifugation at 300 g for 5 min at 4°C, washed twice with ice-cold PBS, and lysed in RIPA buffer for 30 min on ice with intermittent vortexing. Lysates were clarified by centrifugation at 15,000 g for 15 min at 4°C, and supernatants were collected for downstream analysis. Protein concentrations were determined using the BCA Protein Assay Kit (Thermo Fisher Scientific, 23225) according to the manufacturer’s instructions. Equal amounts of total protein were adjusted to the same final volume with lysis buffer, mixed with NuPAGE LDS Sample Buffer (4×; Invitrogen, NP0007) and NuPAGE Reducing Agent (10×; Invitrogen, NP0004), and heated at 70°C for 10 min prior to electrophoresis. Proteins were separated by SDS–PAGE using NuPAGE Bis-Tris gels with MOPS SDS Running Buffer (1×; Invitrogen, NP0001) at 100 – 200 V until the dye front reached the bottom of the gel. Protein standards (Invitrogen, LC5800) were included for molecular weight estimation. Following electrophoresis, proteins were transferred onto PVDF membranes using Tris–glycine transfer buffer (1×; Invitrogen, LC3675) containing 20% methanol at 20 V for 2 h. Membranes were briefly washed with deionized water and stained with Ponceau S solution (Cell Signaling Technology, 59803S) to assess protein transfer and loading. After documentation, membranes were destained with TBST and blocked in 5% non-fat dry milk prepared in TBST for 1 h at room temperature with gentle agitation. Blocked membranes were incubated with primary antibodies diluted in blocking buffer for overnight at 4°C, including MYC (Santa Cruz Biotechnology, sc-40, 1:500; Cell Signaling Technology, 5605T, 1:1000; Invitrogen, MA5-42675, 1:500; Invitrogen, MA1-980-HRP, 1:500; or Abcam, ab32072, 1:1000;), Vinculin (CST, 13901, 1:2000), LC3 (Proteintech, 14600-1-AP, 1:2,000), STING (CST, 50494, 1:2,000).

Membranes were then washed three times with TBST and incubated with appropriate HRP-conjugated secondary antibodies diluted 1:10,000 – 1:20,000 in blocking buffer for 1 h at room temperature. After three additional washes, protein signals were detected using SuperSignal West Pico PLUS chemiluminescent substrate (Thermo Fisher Scientific, 34580) and visualized by chemiluminescence imaging. Band intensities were quantified using ImageJ (NIH). Ponceau S staining or housekeeping proteins were used as loading controls, as indicated.

### Immunofluorescence

Cells were seeded at a density of 5×10^4^ cells per well in 8-well Lab-Tek II CC2 chamber slides (Thermo Scientific) in 250 µL of complete growth medium and allowed to adhere overnight. Cells were transiently transfected as indicated, and immunofluorescence staining was performed 48 h post-transfection. Cells were washed once with PBS and fixed with 4% paraformaldehyde in PBS for 15 min at room temperature. Following fixation, cells were washed three times with PBS and permeabilized using the permeabilization solution in the Image-iT Fixation/Permeabilization Kit (Thermo Fisher Scientific, A5818101) for 15 min at room temperature. Cells were then washed three times with PBS and blocked with the kit-provided blocking solution for 1 h at room temperature. Primary antibody incubation was performed using anti-MYC (Abcam, ab32072), anti-HA antibody (Thermo Fisher Scientific, 26183) diluted 1:500 in blocking solution, followed by overnight incubation at 4°C. Cells were washed three times with PBS and incubated with Alexa Fluor cy3–conjugated goat anti-rabbit IgG (H+L) secondary antibody (Thermo Fisher Scientific, A10520), or Alexa Fluor cy3–conjugated goat anti-mouse IgG (H+L) secondary antibody (Thermo Fisher Scientific, M30010), diluted 1:500 in PBS, for 1 h at room temperature in the dark. After three additional washes with PBS, nuclei were counterstained with DAPI (1 µg/mL in PBS) for 5 min at room temperature in the dark. Following final washes, chambers were mounted with coverslips prior to imaging. Images were acquired at randomly selected fields of view using fluorescence microscopy under identical acquisition settings for all conditions.

Immunofluorescence signals were quantified using ImageJ (NIH). For each cell, nuclear regions of interest (ROIs) were manually delineated based on DAPI staining, while cytoplasmic ROIs were defined as the total cell area minus the nuclear region. Integrated fluorescence intensity and area were measured for each ROI. Background fluorescence was determined from an adjacent cell-free region and subtracted using the corrected total cell fluorescence (CTCF) formula: CTCF = Integrated Density − (Area × Mean Background Fluorescence).

### RNAScope

Cells were seeded one day prior to transfection in 8-well Lab-Tek II CC2 chamber slides at a density sufficient to achieve 70-80% confluence at the time of fixation and allowed to adhere firmly. Cells were transfected using the same procedure as described above and a final ASO concentration of 100 nM. After 48 h transfection, cells were washed twice with PBS and fixed with 10% neutral buffered formalin (Sigma-Aldrich, HT501128) for 30 min at room temperature. Following fixation, cells were rinsed three times with PBS and processed immediately for RNAScope analysis. RNAScope multiplex fluorescent in situ hybridization was performed using the RNAScope Multiplex Fluorescent Reagent Kit v2 (ACD Bio, 323110) according to the manufacturer’s instructions with minor modifications. All hybridization steps were carried out using the HybEZ II Hybridization System at 40°C unless otherwise specified. Wash buffer was prepared by diluting the 10× RNAScope wash buffer to 1× with distilled water. Fixed cells were treated with RNAScope hydrogen peroxide for 10 min at room temperature to quench endogenous peroxidase activity, followed by two washes with distilled water. Cells were then permeabilized by incubation with diluted Protease III (1:15 in PBS) for 10 min at room temperature. Cells were washed three times with PBS prior to probe hybridization. Target probe hybridization was performed using the human *MYC* probe (Hs-MYC-C3), diluted according to the manufacturer’s recommendations, and incubated for 2 h at 40°C. Excess probe was removed, and cells were washed three times with 1× RNAScope wash buffer. Signal amplification was carried out sequentially using RNAScope Multiplex FL v2 Amp 1 and Amp 2 reagents for 30 min each, followed by Amp 3 for 15 min, with two washes in wash buffer between each amplification step. For signal detection, HRP-C3 was applied for 15 min at 40°C, followed by washing and development with Opal 690 fluorophore (Akoya Biosciences, FP1497001KT), which substitutes for TSA Plus Cyanine 5. Opal 690 was diluted 1:1500 in RNAScope Multiplex TSA buffer and incubated for 30 min at 40°C. Following fluorophore development, HRP activity was quenched using the RNAScope HRP blocker for 15 min at 40°C, followed by washing. Nuclei were counterstained with DAPI for 30 s at room temperature. Cells were washed with PBS and chambers were mounted with coverslips prior to imaging. Fluorescence images were acquired at randomly selected fields of view using identical acquisition settings across conditions.

RNAScope signals were quantified using ImageJ (NIH). For each field of view, individual RNA puncta located in the cytoplasm were identified, and nuclear boundaries were manually or semi-automatically delineated based on DAPI staining. Only RNA puncta outside the nuclear boundary were included for analysis. The distance from each cytoplasmic RNA punctum to the nearest nuclear boundary was measured using ImageJ measurement tools.

### Apoptosis analysis by Annexin V/7-AAD staining

Cells were seeded in 6-well plates at a density of 1.0 × 10^5^ cells per well and transiently transfected as indicated. 72 h after transfection, cells were harvested for apoptosis analysis. Cells were washed twice with ice-cold Cell Staining Buffer (BioLegend, 420201) and resuspended in Annexin V Binding Buffer (BioLegend, 422201). A total of 100 µL of the cell suspension was transferred to 1.5 mL tubes for staining. Fluorochrome-conjugated Annexin V (BioLegend, 640930), diluted 1:10 in Annexin V Binding Buffer, was added at 5 µL per sample, followed by the addition of 7-aminoactinomycin D (7-AAD) according to the manufacturer’s instructions. Samples were gently mixed and incubated for 15 min at room temperature in the dark. Following incubation, 400 µL of Annexin V Binding Buffer was added to each sample, and cells were analyzed immediately by flow cytometry. Data were analyzed using standard flow cytometry analysis software (FCS Express).

### Single-cell cloning and genotyping

D11-SpRY-Cas9 cells were seeded at 2.0×10^6^ cells per T25 non–tissue-culture-treated flask in 5 mL complete growth medium. Lentiviral particles encoding sg723 or sgCtrl, packaged as described above, were added directly to the cultures (20 µL virus per flask) together with polybrene (8 µg/mL). Culture medium was replaced 24 h post-infection. Cells were collected by centrifugation at 200 g for 5 min, washed twice with fresh medium (200 g for 5 min each wash), resuspended in complete medium, and subsequently expanded and passaged as needed. To assess infection efficiency and ensure appropriate conditions for single-cell cloning, the percentage of mCherry-positive (sgRNA-positive) cells was quantified by flow cytometry. mCherry-positive cells infected with sgCtrl were collected and expanded as polyclonal control populations. For clonal isolation, cells infected with sg723 were single-cell sorted into 96-well plates (one cell per well) using a Sony MA900 cell sorter and cultured under standard conditions. Individual clones were expanded, and genomic DNA was isolated using the Macherey-Nagel NucleoSpin Tissue kit (Macherey-Nagel, 740952.50) according to the manufacturer’s instructions. The targeted genomic locus was amplified using Q5 High-Fidelity 2× Master Mix (New England Biolabs, M0492S) with primers listed in Supplementary Table 4. PCR reactions (50 µL total volume) contained 25 µL Q5 High-Fidelity 2× Master Mix, 2.5 µL of each primer (10 µM; final concentration 0.5 µM), template genomic DNA (<1,000 ng), and nuclease-free water to volume. Thermocycling conditions were as follows: initial denaturation at 98°C for 30 s; 40 cycles of 98°C for 10 s, 68°C for 30 s, and 72°C for 90 s; followed by a final extension at 72°C for 2 min and a hold at 4–10°C. PCR products were resolved by agarose gel electrophoresis, purified, and subjected to amplicon sequencing using the Plasmidsaurus sequencing service. Sequencing data were analyzed as described above.

### RNA-seq

MDA-MB-231 cells were cultured under standard conditions. Cells were seeded in 6-well plates at 1.0×10⁵ cells per well for the negative control conditions (ASO-ctrl-1, ASO-ctrl-2, and ASO-ctrl-3) and at 3.0×10⁵ cells per well for the ASO-treated conditions (ASO-1, ASO-2, and ASO-3). Cells were transfected with the indicated ASOs at a final concentration of 100 nM. Culture medium was refreshed 48 h post-transfection. Cells were collected at <80% confluency. MDA-MB-231 treated cells were harvested at 48 h and 72 h post-transfection. At each time point, cells were lysed directly in 350 µL lysis buffer (LBP), and lysates were immediately stored at −80°C until RNA isolation.

MDA-MB-231 parental cells and D11-SpRY-Cas9 cells were collected without any treatment. cells were lysed directly in 350 µL lysis buffer (LBP), and lysates were immediately stored at −80°C until RNA isolation.

HEK293T cells were cultured under standard conditions. Cells were seeded in 6-well plates at 1.0×10⁵ cells per well. Cells were transfected with the indicated plasmids at a final concentration of 2500 ng. Culture medium was refreshed 48 h post-transfection. Cells were harvested at 72 h post-transfection. Cells were lysed directly in 350 µL lysis buffer (LBP), and lysates were immediately stored at −80°C until RNA isolation.

Total RNA was extracted using the MACHEREY-NAGEL NucleoSpin RNA Plus kit (Macherey-Nagel, 740984.250) according to the manufacturer’s instructions. RNA purity was assessed by NanoDrop spectrophotometry, only samples with A_260_/A_280_ ratios of approximately 2.0 and A_260_/A_230_ ratios greater than 2.0 were used. RNA integrity was evaluated using the Agilent TapeStation, and only samples with RNA integrity number equivalent (RINe) value greater than 9.5 were used for RNA sequencing. RNA-sequencing libraries were prepared from quality-controlled total RNA using poly(A) selection to enrich for mRNA. Libraries were constructed following standard bulk RNA-sequencing protocols and sequenced on the AVITI sequencing platform to generate transcriptomic data.

### sgRNA phenotype score calculation

The number of reads per sgRNA was computed with MAGeCK^26^ (version 0.5.9.5) *count* subcommand and normalized with negative control sgRNAs (i.e. targeting AAVS1, non-targeting, or random). sgRNA-level log2 fold change for each sample was computed with MAGeCK’s *test* subcommand using normalized count as input with parameters *--norm-method none --adjust-method fdr --control-gene control_gene_list.txt*. Day 8 and day 20 samples were compared to day 0 to calculate log2 fold changes. The two day 0 samples generated using high- and low-MOI infections produced highly similar results. We therefore used the high-MOI day 0 sample for all subsequent analyses, as higher MOI is expected to better preserve library complexity. Log2 fold change was further normalized for each sample by the number of days in culture to allow comparison. sgRNA-level phenotype score was calculated as the average of sgRNA-level per day log2 fold change across four samples.

### Base-pair phenotype score calculation

To mitigate phenotype-independent variations across sgRNAs, base-pair-level log2 fold change was calculated by averaging the sgRNA-level log2 fold change of the eight sgRNAs (four per strand) predicted to cut within ±1 bp of each base-pair. We leveraged MAGeCK gene-level summarization to achieve this. First, a pseudo MAGeCK count table was generated from normalized sgRNA count table with a customized script so that each sgRNA was duplicated four times and each duplicate’s “Gene” column was mapped to a base-pair coordinate within ±1 bp of its cutting site. Next, base-pair-level log2 fold change was computed with MAGeCK *test* subcommand using pseudo count table as input and high MOI day 0 sample as baseline with parameters *--norm-method none --gene-test-fdr-threshold 0.01 --adjust-method fdr --gene-lfc-method mean --sort-criteria neg --control-sgrna control_sgrna_list.txt*. With the modified pseudo count table, base-pair-level log2 fold change as the average of eight sgRNAs would be outputted as gene-level log2 fold change by MAGeCK test. One advantage of leveraging MAGeCK is that it uses robust ranking aggregation (RRA) algorithm to identify significantly regulated genes (base-pairs in our scenario) by comparing the skew in sgRNA rankings of a gene to the uniform null model and prioritizing genes whose sgRNA rankings are consistently higher than expected. In our case, base-pairs whose eight sgRNAs are consistently showing stronger phenotypes than negative control sgRNAs would be prioritized. Base-pair-level log2 fold change was further normalized for each sample by the number of days in culture to allow comparison. Base-pair-level phenotype score was calculated as the average base-pair-level per day log2 fold change across four samples. The median score of control sgRNAs with no off-targets in human genome when allowing at most one mismatch (474 control sgRNAs), which were randomly grouped for 1000 iterations with eight sgRNAs in a group to match base-pair-level averaging strategy, was subtracted from base-pair-level phenotype scores. A base-pair was considered significant if it showed an FDR < 0.05 in at least two samples with at least two strong sgRNAs in each and exhibited a consistent log2 fold change directional trend in the remaining samples. Only 13 out of 22,058 MYC locus targeting sgRNAs and no negative control sgRNAs have low sequencing coverage in baseline sample (raw read count lower than 100). They were not removed given their low proportion in the entire sgRNA library and that their effects were mitigated as the base-pair-level phenotype score was calculated as the average of eight sgRNAs. Base-pairs within the GFP region were not considered when calculating the base-pair phenotype score.

### Gene and feature annotations

Human reference genome assembly hg38 was used throughout all analyses. *MYC* transcript ENST00000377970.6 was used throughout all analyses since it is the main *MYC* isoform expressed in the D11 screening cell line as shown by D11 RNA-Seq data. The possible mutation outcomes of a given base-pair were predicted by Variant Effect Predictor (Ensembl version 112)^76^ using ENST00000377970.6 as the reference *MYC* transcript. Since the category (i.e. coding, UTRs, etc.) of mutation outcomes in mutagenesis screening largely depends on the position of the cutting site instead of the exact indels, which are unknown at each position, it was assumed that an A was inserted for every cutting site when submitting to Variant Effect Predictor. Base-pairs were further categorized into “upstream_flanking”, “5’UTR, “intron”, “intron_splice”, “coding”, “3’UTR”, and “downstream_flanking” based on their most damaging mutation outcomes (e.g. base-pairs around splice sites were categorized as “intron_splice” instead of “introns”, unless otherwise specified. Variant Effect Predictor defines variant within 1-8 bases at the 5’ end of an intron or 1-17 bases at the 3’ end of an intron as splice variant). Base-pairs within 1-kb region upstream of the transcription start site of transcript ENST00000377970.6 were further categorized as “promoter” unless otherwise specified. Repeat annotation was retrieved from RepeatMasker^77^. A sgRNA was considered falling into repeat regions if the 20-nt sgRNA entirely overlapped with repeats regardless of the strand. A base pair would be labeled as repeats if it was within repeat regions regardless of the strand. All base pairs within repeats were labeled or removed in analyses unless otherwise noted. ENCODE3 candidate *cis*-regulatory elements (cCREs) annotation combined from all cell types^33^ was downloaded from UCSC Genome Browser. Base-pairs were categorized into falling within elements with promoter-like signature (prom), proximal enhancer-like signature (enhP), distal enhancer-like signature (enhD), or no cCRE signature.

### Off-target and target efficiency prediction

Potential off-targets of sg723 were searched by CasOFFinder^44^ web server with parameters PAM type “SpRY Cas9 from Streptococcus pyogenes: 5’-NNN-3” and target genome “Homo sapiens (GRCh38/hg38)”. Off-target sites with mismatch numbers no more than two and no DNA bulges or RNA bulges were further mapped to genes according to GENCODE human release 47 annotations. Off-target indels at these sites were experimentally measured by amplicon sequencing using cells expressing corresponding sgRNA. Potential off-targets of all *MYC*-targeting and negative control sgRNAs were searched by mapping sgRNA sequences to human genome GRCh38.p14 with bowtie^78^ (version 1.2.3,) allowing one mismatch (-v 1 –all). Bam outputs from bowtie were processed with pysam (version 0.21.0) to obtain sgRNA-level off-target counts. On-target efficiency of sgRNA was predicted with CRISPROn^30^ (version 1.0), DeepSpCas9^31^, and CRISPick^29^. Given the PAM motif constraints of all three prediction tools, only sgRNAs with NGG PAM motif were considered for on-target efficiency prediction. To ensure that the observed growth phenotype differences reflected on-target efficiency rather than variation in sgRNA target regions, only sgRNAs with NGG PAM motif and targeting at the coding region in exon 2 were considered in the comparison between predicted on-target efficiency and sgRNA phenotype score.

### Conservation – function correlation

Whole-genome phastCons scores^79^ derived from multiple alignments of 99 vertebrate genomes to the human genome was downloaded from UCSC Genome Browser. Coding regions on the whole genome level were defined as CDS regions of MANE Select transcripts^80^ (MANE release 1.4). *MYC* coding regions were specifically defined as the coding regions in transcript ENST00000377970.6. Per-base-pair JARVIS^40^ score on 10-kb *MYC* locus in hg38 was downloaded from UCSC Genome Browser. To match the averaging strategy of the base-pair phenotype score, per-base-pair phastCons scores were smoothed using a 5-bp sliding window, in which the score at each base-pair was replaced by the average of scores within ±2 bp of that position. JARVIS scores were smoothed in the same way. *RPS19* SpRY-mediated screening data was retrieved from supplementary table 2 and supplementary table 3 in Yao *et al*^41^. Two supplementary tables were merged together, and average log2(CRISPR score)2 as provided in tables was taken as phenotype score. Since the provided CRISPR score was calculated by averaging the sgRNA scores of five consecutive sgRNAs centered at the indicated sgRNA, *RPS19* phastCons scores were smoothed in the same manner as *MYC* phastCons scores.

### 3’ UTR conservation and mRNA half-life

The aggregated half-life data of human genes was downloaded from Agarwal *et al*^81^. 3’UTR coordinates of MANE Select protein-coding transcripts were retrieved from UCSC Genome Browser Table Browser (MANE release 1.4). Whole-genome per-base-pair phyloP^82^ score for multiple alignments of 99 vertebrate genomes to the human genome was downloaded from UCSC Genome Browser. Highly conserved base-pairs in 3’UTR were defined with the method from Luo et al^83^. A base-pair was considered highly conserved if its phyloP score is no less than 2 and at least four consecutive base-pairs had phyloP scores above this cutoff.

### Visualization with UCSC Genome Browser

Screening signal along *MYC* locus was visualized using customized plotting scripts in combination with UCSC Genome Browser^84^. Standard tracks were provided directly by UCSC Genome Browser, including GENCODE v49 annotation, phastCons conservation score, GC percent in 5-base windows, DNase signals, and ENCODE3 candidate *cis*-regulatory elements (cCREs) annotation. RNA-Seq coverage track was generated from the screening cell line D11 RNA-Seq data (method specified in RNA-Seq analysis).

### ASO RNA-Seq analysis

RNA-Seq fastq files were processed with fastp^85^ (version 0.23.2) using default parameters to filter out low-quality reads, trim low-quality bases and remove adapters if needed. Reads were mapped to genome assembly hg38 (GRCh38.p14) with STAR^86^ (version 2.7.10b) using parameter --outFilterMultimapNmax 1 (unique mappers only). Reads were quantified with featureCounts^87^ (version 2.0.1) using parameters -s 2 -p and GENCODE human release 47 annotations. Transcripts Per Million (TPM) was calculated for each gene based on the raw count table and gene length. Coverage track was generated from mapping bam files with deepTools^88^ bamCoverage (version 3.5.4) using parameters --binSize 5 --normalizeUsing None –centerReads. Differential expression analysis was carried out with DESeq2 (version 1.46.0)^46^. Samples 48-hour and 72-hour post ASO treatment were compared to control samples of corresponding time points. Genes were ranked according to DESeq2 “stat” column output and gene set enrichment analysis (GSEA) was carried out with clusterProfiler (version 4.14.6)^89^. Gene sets were retrieved from the Molecular Signatures Database (MSigDB).

### Statistics and Reproducibility

In general, all experiments were performed with at least three independent biological replicates, except for the HF-SpRY-Cas9 screen, which was performed twice starting from independent HF-SpRY-Cas9 stable cell line generations. Statistical analyses were conducted using GraphPad Prism software (version 10.4), except where specific statistical methods are indicated in the figure legends or described above. Comparisons between two groups were analyzed using a two-tailed Student’s t-test, while one-way ANOVA was used for comparisons involving more than two groups. Representative data were derived from at least three independent replicates. Data are presented as mean ± standard deviation (SD). Statistical significance is indicated as follows: ns, not significant, *: p<0.05, **: p<0.01, ***: p<0.001, ****: p<0.0001.

---

**Figure S1:**
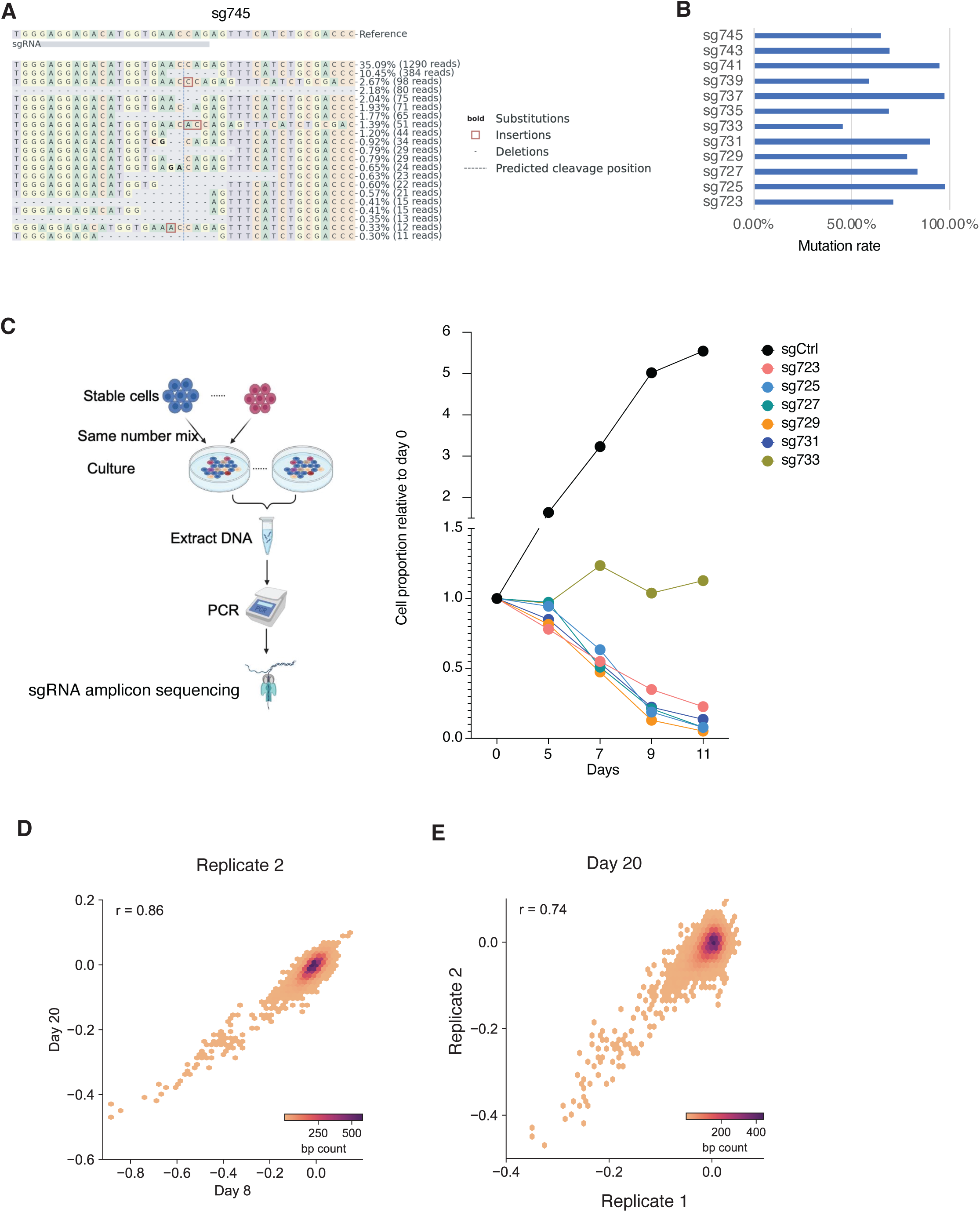
Validation and quality control of the saturation mutagenesis screen. **A**. Representative mutational patterns uncovered by amplicon sequencing for one sgRNA (sg745). **B**. Mutation rate for all 12 sgRNAs tested by amplicon sequencing, calculated as 100% – WT%. **C**. Competitive growth assay for 6 of the 12 sgRNAs. Data were normalized to Day 0. **D**. Scatter plot of base-pair level phenotype scores between day 8 and day 20 samples (replicate 2). **E**. Scatter plot of base-pair level phenotype scores between replicate 1 and replicate 2 (day 20 samples).

**Figure S2:**
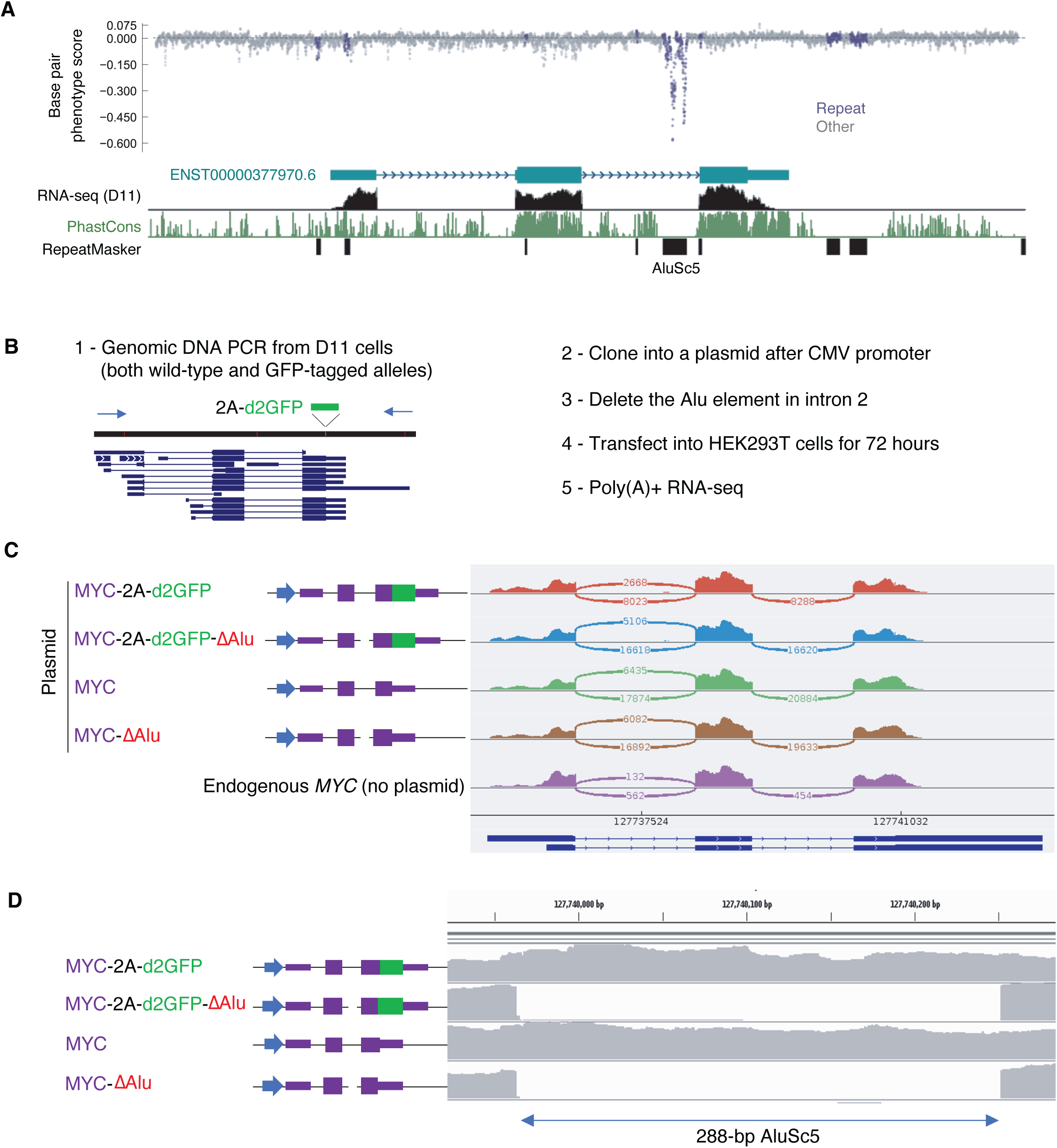
An Alu repeat element within *MYC* intron 2. **A**. An AluSc5 repeat element within *MYC* intron 2 shows the strongest deleterious effect in the CRISPR saturation mutagenesis screen. **B**. Experimental strategy to assess the role of the Alu element in *MYC* expression by cloning the *MYC* gene into plasmids with or without deletion of the Alu element. D11 cells harbor one wild-type *MYC* allele and one GFP-tagged allele; both alleles were cloned and tested. **C**. Schematic of plasmid constructs containing the *MYC* gene with or without GFP and with or without the Alu element. RNA-seq read coverage and splice junctions are shown on the right. Non-transfected HEK293T cells expressing endogenous *MYC* are included as a reference. The lower read coverage observed in the first sample (MYC-2A-d2GFP) likely reflects variation in plasmid concentration or transfection efficiency despite attempts to match conditions across samples. **D**. RNA-seq read coverage across the Alu element confirms successful deletion of the Alu sequence in the corresponding constructs.

**Figure S3.**
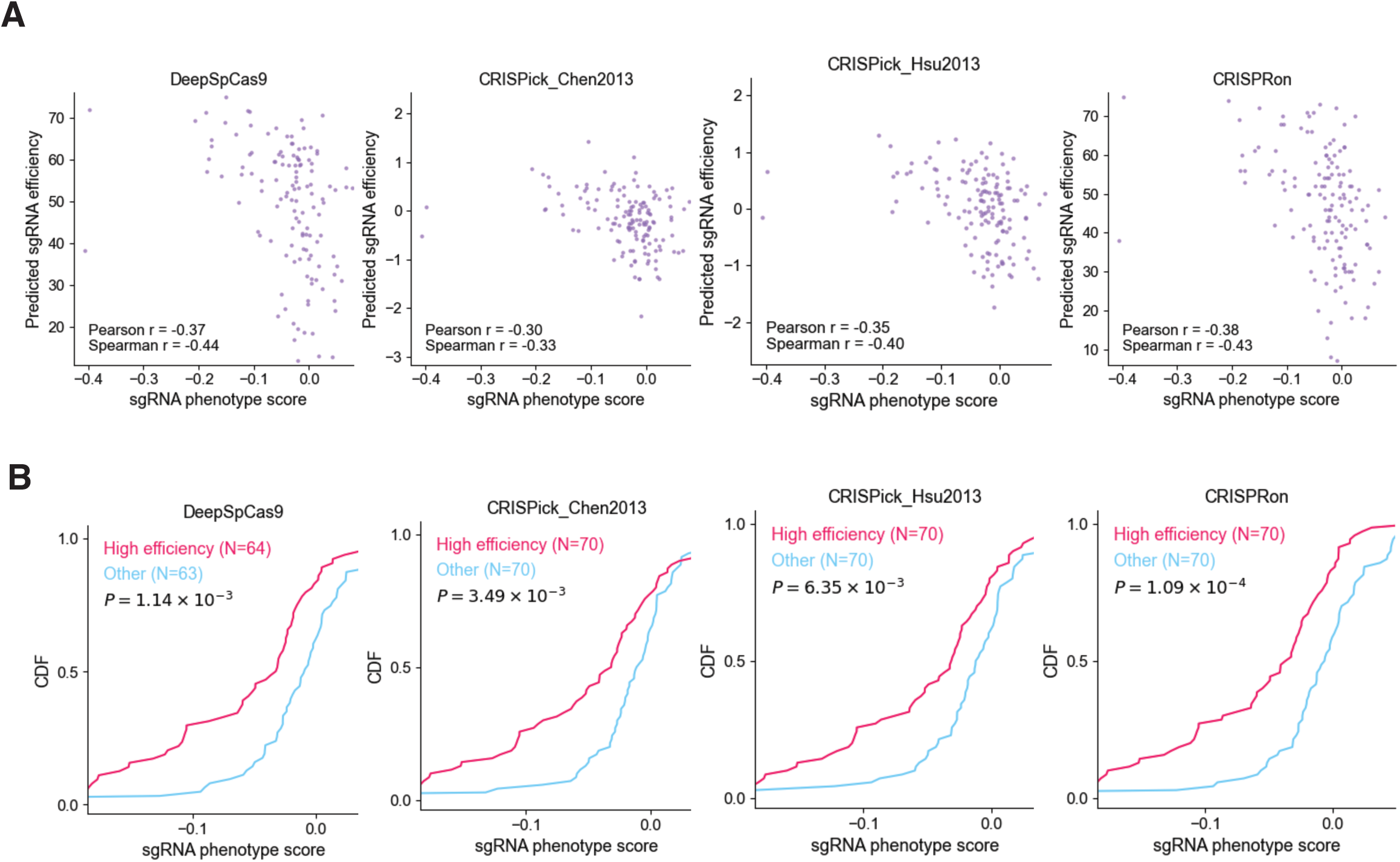
Concordance between predicted sgRNA efficiency and observed phenotype scores. **A**. Scatter plots showing expected negative correlations between observed sgRNA phenotype scores and sgRNA efficiency predicted by four computational tools. Only sgRNAs with NGG PAM motif and targeting at the coding region in exon 2 were considered. **B**. Cumulative distribution function (CDF) plots comparing sgRNA phenotype scores between sgRNAs with high (top half) and low (bottom half) predicted efficiency. Kolmogorov–Smirnov test was used to compute the P values.

**Figure S4.**
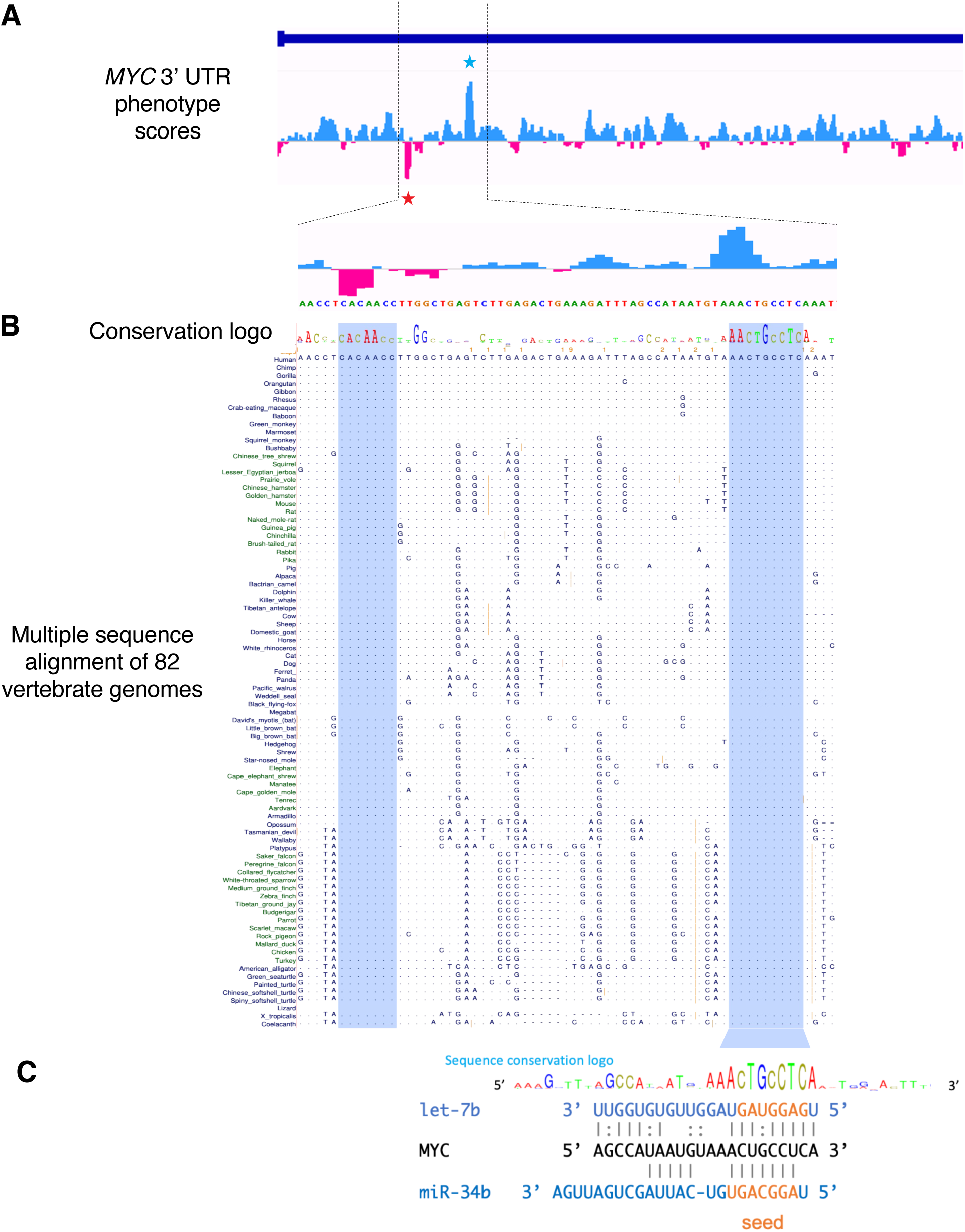
The strongest positive and negative phenotypes in *MYC* 3’ UTR map to two adjacent ultraconserved elements. **A**. Base-pair phenotype scores in *MYC* 3’ UTR, with zoom-in to the region that contains the strongest positive and negative phenotype scores. Blue and red stars highlight 3’ UTR base pairs with the strongest positive and negative phenotype scores, respectively. **B**. PhyloP sequence conservation logo (UCSC genome browser) and multiple sequence alignment. The two ultraconserved element are highlighted in blue, perfectly conserved across 82 genomes. **C**. One of the ultraconserved element contains overlapping seed match sites to two tumor suppressor microRNAs, let-7b and miR-34b.

**Figure S5.**
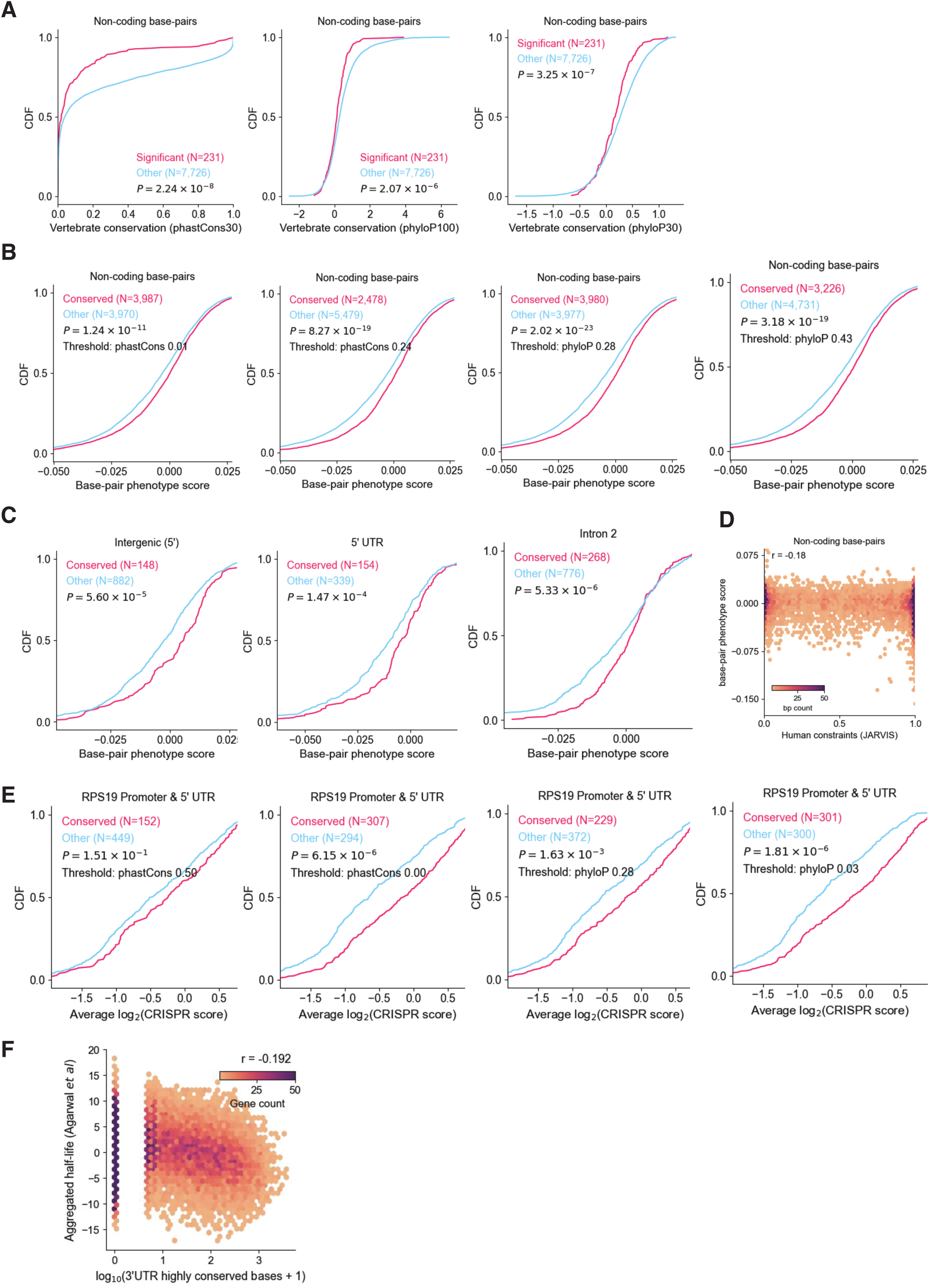
A paradox between conservation and function. **A**. Same as Fig. 3A but using phastCons calculated from 30 mammals (left), phyloP scores from 100 vertebrate genomes (center) or 30 mammals (right). **B**. Same as Fig. 3B but using different thresholds of phastCons scores or phyloP scores (median and mean values of non-coding base-pairs). **C**. Same as Fig. 3C for the intergenic region upstream of the promoter, 5’ UTR, and intron 2. Other regions are not significant. **D**. Weak correlation between JARVIS human constraints and base-pair phenotype scores. **E**. Same as in B using RPS19 promoter and 5’ UTR saturation mutagenesis data from Yao *et al*. **F**. The number of conserved base-pairs in 3’ UTRs is negatively correlated with average half-lives of mRNAs. P value = 5.6x10^-118^. Kolmogorov–Smirnov test was used to compute all the P values.

**Figure S6.**
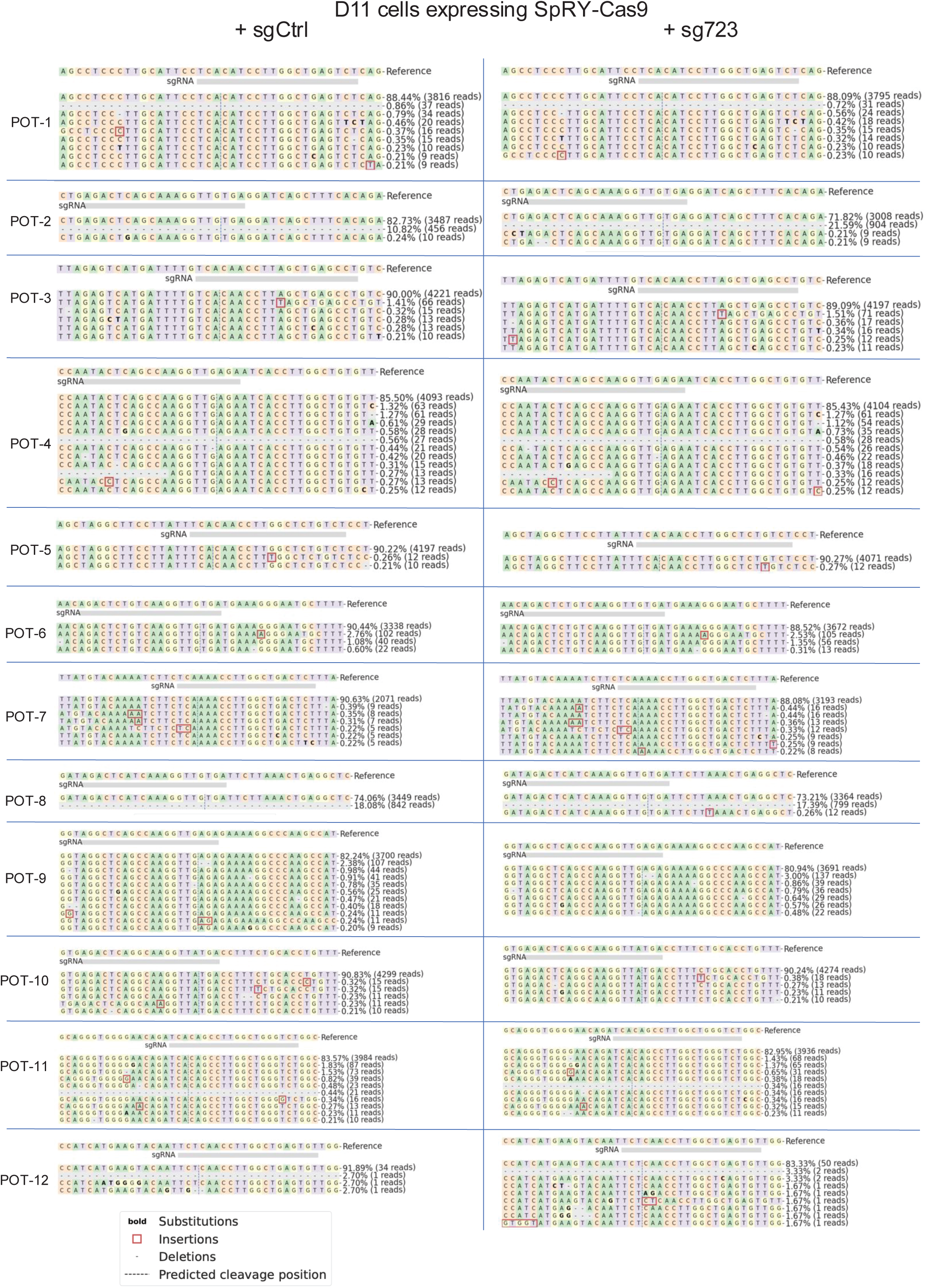
Lack of mutations at predicted off-target sites for sg723. D11 cells stably expressing SpRY-Cas9 were further transduced with either sg723 or sgCtrl. Mutations at the twelve predicted off-targets (POT) were assayed using amplicon sequencing. No significant mutations were detected at these sites. Some low frequency mutations were detected in both sg723 and sgCtrl samples, likely resulting from sequencing errors.

**Figure S7.**
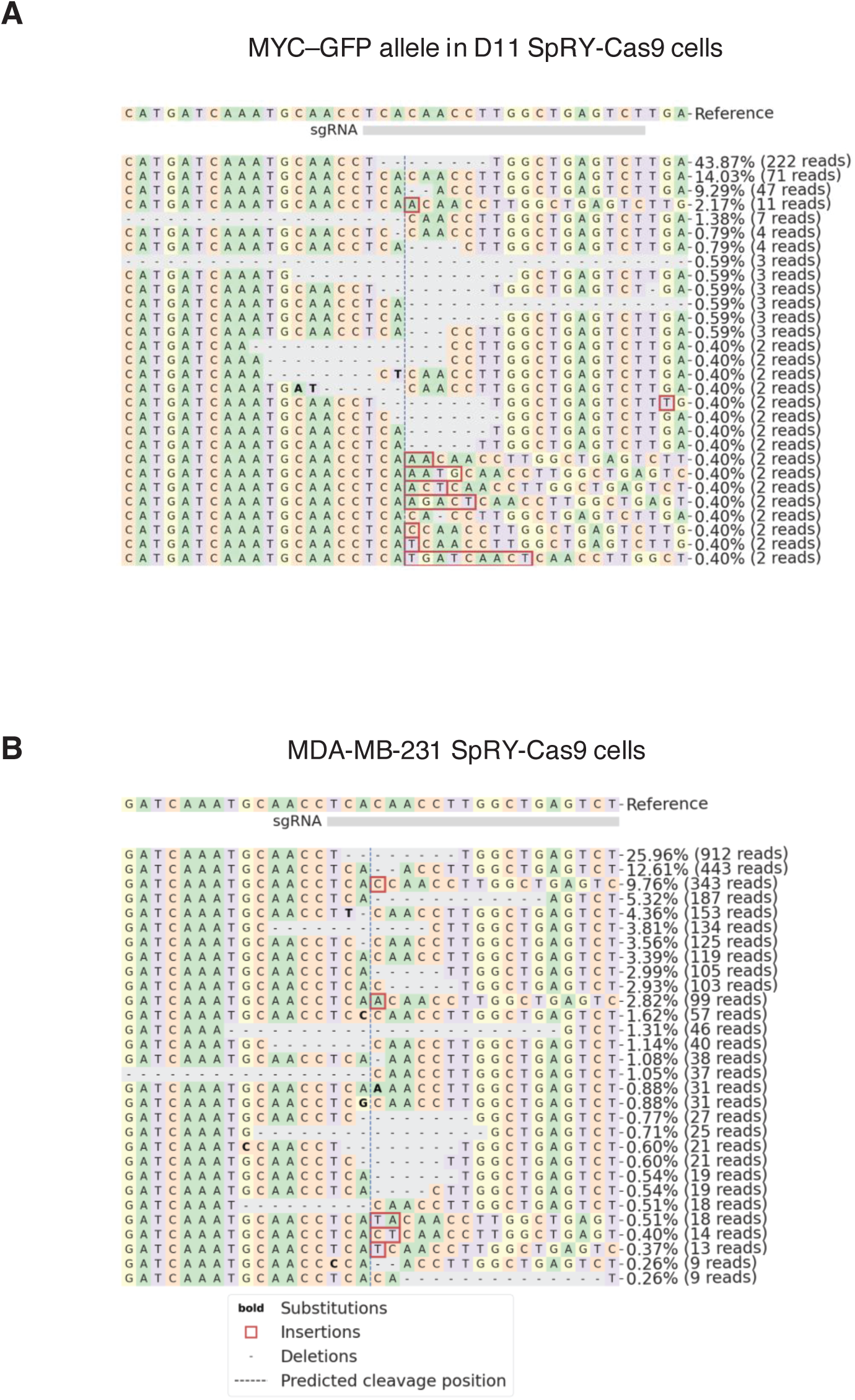
Mutational patterns of sg723. **A**. Amplicon sequencing of the GFP-tagged MYC allele in D11 cells targeted with sg723, related to Fig. 4A. **B**. Amplicon sequencing of sg723 target sites in MDA-MB-231 cells.

**Figure S8.**
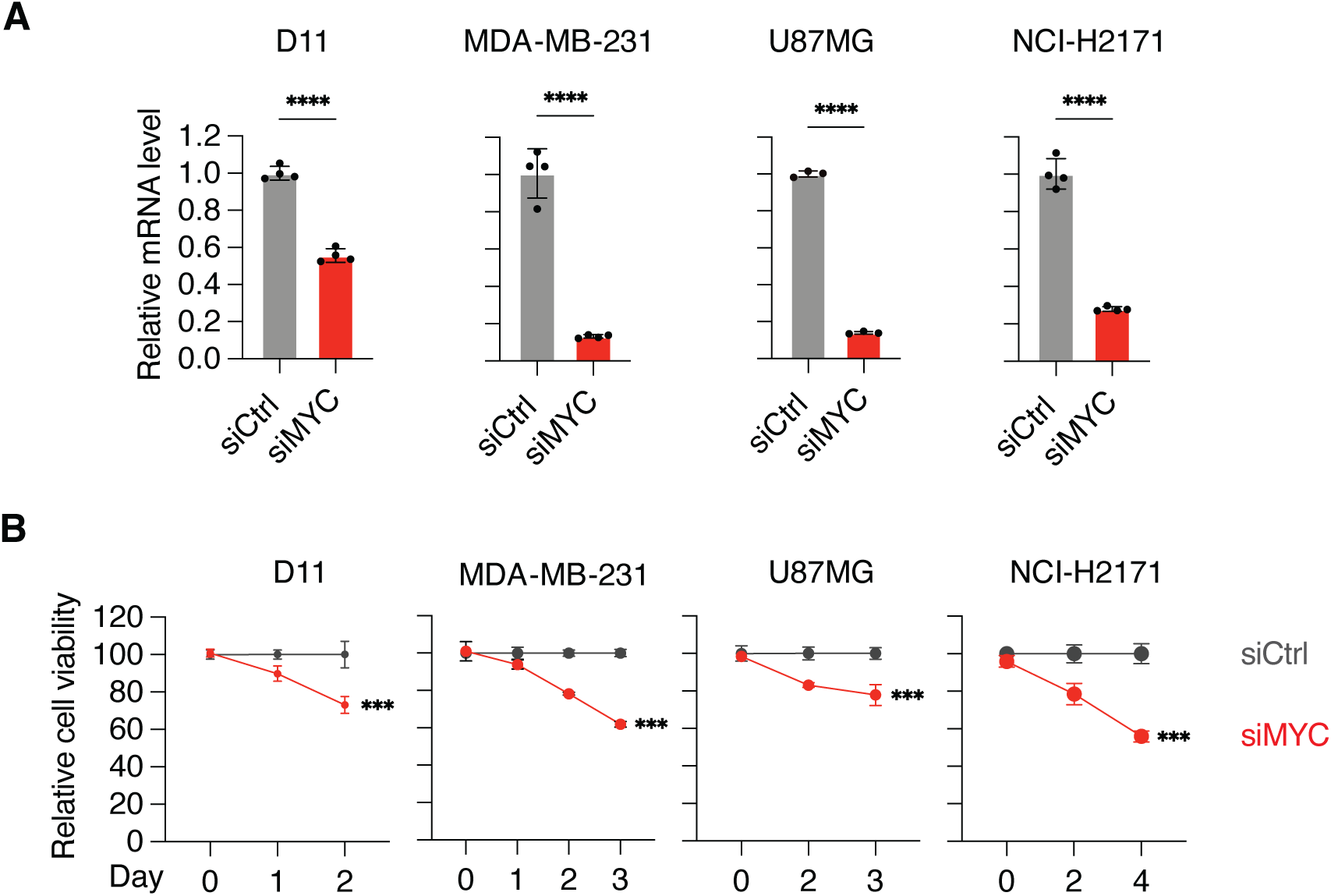
MYC-dependent cell growth. **A**. siRNA-mediated knockdown of *MYC* mRNA in indicated cell lines. Quantified with RT-qPCR using *ACTB* as internal control. Data were normalized to siCtrl. **B**. Relative cell viability of indicated cell lines with siRNA-mediated knockdown of *MYC*. Data were normalized to siCtrl. ***: p<0.001, ****: p<0.0001.

**Figure S9.**
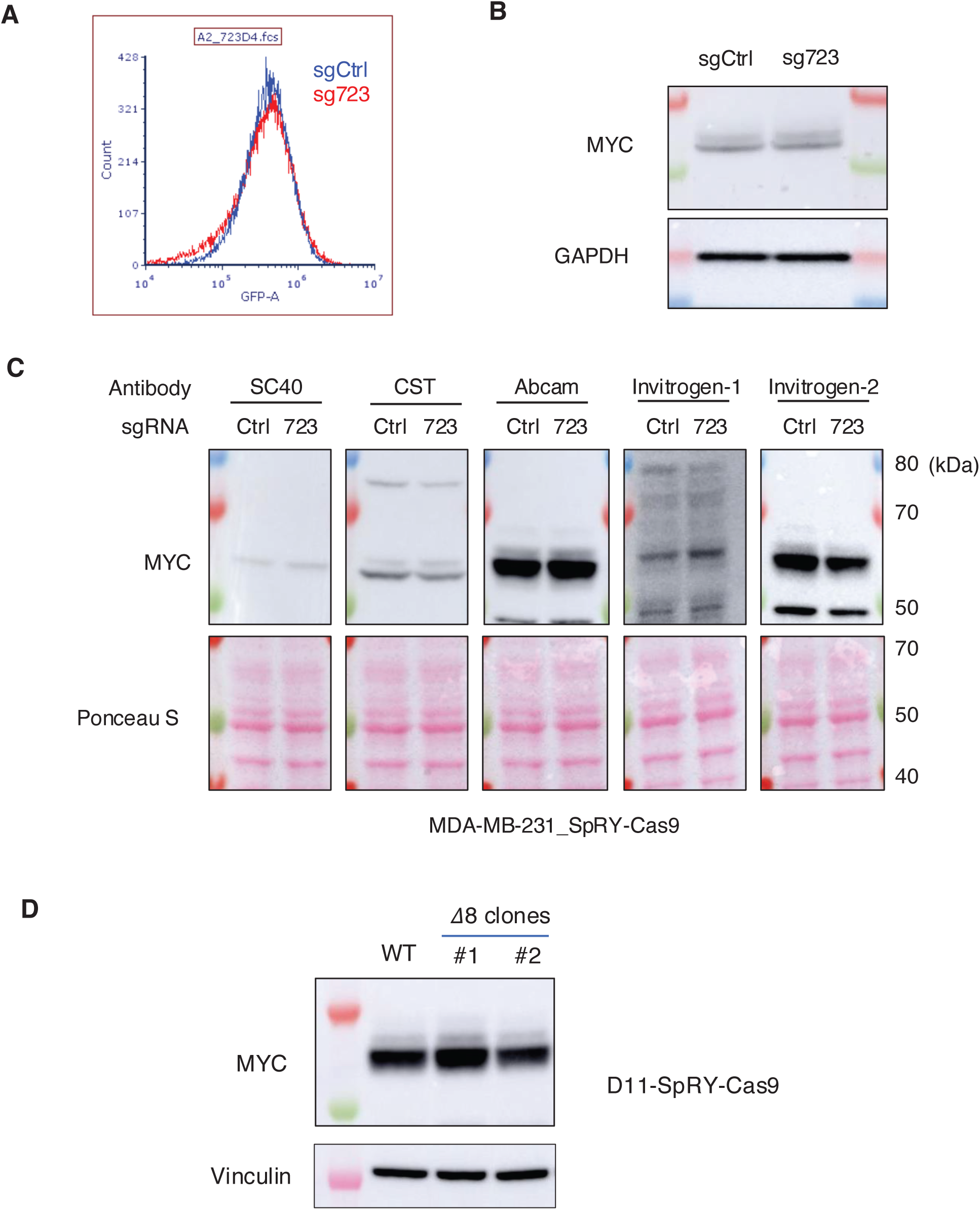
Lack of change in MYC abundance. **A**. Flow cytometry quantification of GFP signal in SpRY-Cas9 D11 cells transduced with either sgCtrl or sg723. **B**. Western blotting of MYC in MDA-MB-231 cells transduced with sgCtrl or sg723. **C**. No significant change of MYC protein abundance in MDA-MB-231 cells when detected using five commercially available antibodies. Ponceau S was used as loading control. **D**. No significant change of MYC protein abundance in D11 clonal cells with the 3’ UTR element disrupted (𝛥8).

**Figure S10.**
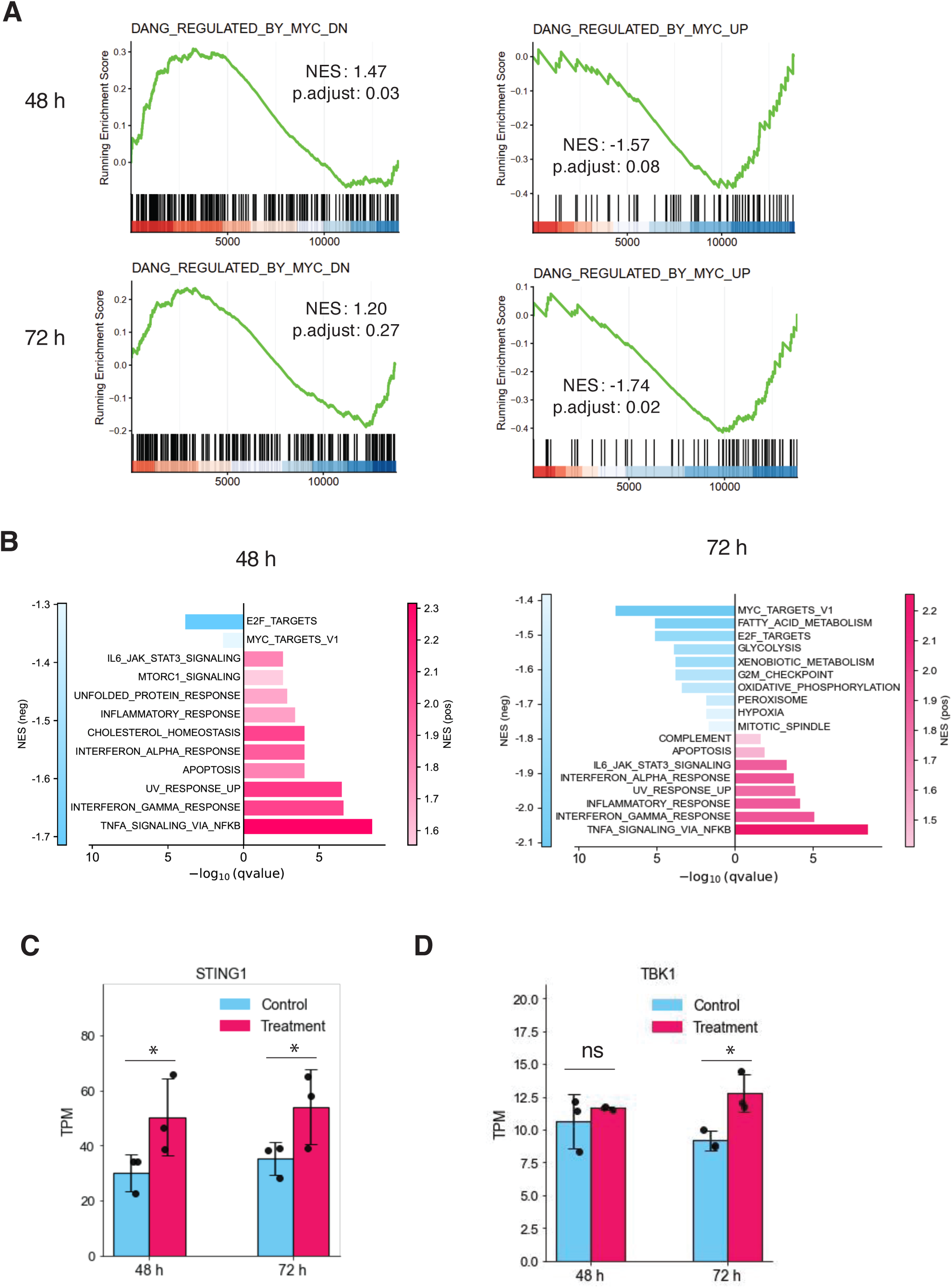
Genes and pathways affected by MYC ASOs. **A**. Gene set enrichment analysis (GSEA) plots for MSigDB MYC-downregulated gene set and MYC-upregulated gene set in 48 or 72 h after ASO treatment. **B**. MSigDB hallmark gene sets significantly enriched in down- or up-regulated genes (ASO treated vs ASO control). **C**. Expression change of STING1. **D**. Expression change of TBK1. TPM: Transcripts per million. Significance reflects adjusted p-values from DESeq2 differential expression analysis performed on raw counts.

**Figure S11.**
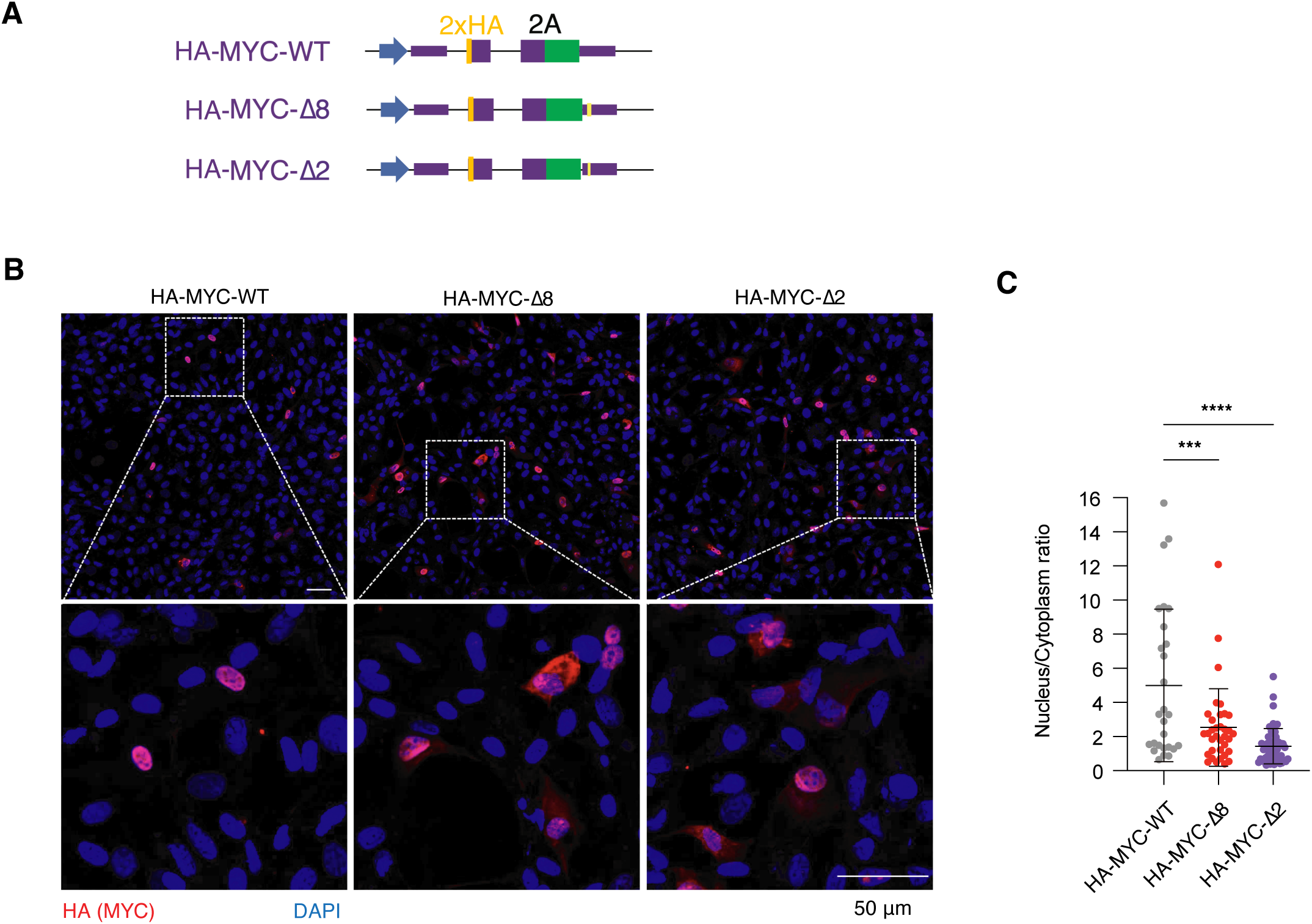
Deleting the ultraconserved 3’ UTR element alters MYC protein localization. **A**. HA-tagged MYC was expressed from a plasmid containing the full-length *MYC* gene, including introns and UTRs. The HA-MYC-𝛥8 and -𝛥2 variants delete 8-nt and 2-nt from the element, respectively. Plasmids were transiently transfected into U87MG cells for 48 h, and MYC localization was assessed by HA-tag immunofluorescence staining. **B**. Representative immunofluorescence images. **C**. Quantification of the nucleus/cytoplasm ratio as in Fig. 6A/B. MYC-WT: N=27; MYC-𝛥8: N=35; MYC-𝛥2: N=55. ***: p<0.001, ****: p<0.0001.

